# PCNA activates the MutLγ endonuclease to promote meiotic crossing over

**DOI:** 10.1101/2020.02.12.946020

**Authors:** Dhananjaya S. Kulkarni, Shannon Owens, Masayoshi Honda, Masaru Ito, Ye Yang, Mary W. Corrigan, Lan Chen, Aric L. Quan, Neil Hunter

## Abstract

During meiosis, crossover recombination connects homologous chromosomes to direct their accurate segregation^1^. Defects in crossing over cause infertility, miscarriage and congenital disease. Accordingly, each pair of chromosomes attains at least one crossover through processes that designate and then implement crossing over with high efficiency^2^. At the DNA level, crossing over is implemented through the formation and biased resolution of double-Holliday Junction intermediates^3–5^. A central tenet of crossover resolution is that the two Holliday junctions are resolved in opposite planes by targeting nuclease incisions to specific DNA strands^6^. Although the endonuclease activity of the MutLγ complex has been implicated in crossover-biased resolution^7–12^, the mechanisms that activate and direct strand-specific cleavage remain unknown. Here we show that the sliding clamp, PCNA, is important for crossover-biased resolution. *In vitro* assays with human enzymes show that hPCNA and its loader hRFC are sufficient to activate the hMutLγ endonuclease under physiological conditions. In this context, the hMutLγ endonuclease is further stimulated by a co-dependent activity of the pro-crossover factors hEXO1 and hMutSγ, the latter of which binds Holliday junctions^13^. hMutLγ also specifically binds a variety of branched DNAs, including Holliday junctions, but canonical resolvase activity is not observed implying that the endonuclease incises adjacent to junction branch points to effect resolution. *In vivo*, we show that budding yeast RFC facilitates MutLγ-dependent crossing over. Furthermore, PCNA localizes to prospective crossover sites along synapsed chromosomes. These data highlight similarities between crossover-resolution and DNA mismatch repair^14–16^ and evoke a novel model for crossover-specific dHJ resolution during meiosis.

## Main

Meiotic recombination is initiated by programmed DNA double-strand breaks (DSBs) and proceeds via homologous pairing and DNA strand-exchange to form joint-molecule (JM) intermediates^1^. Regulatory processes designate a subset of events to become crossovers, and at these sites nascent JMs are matured into double-Holliday junctions (dHJs). Through poorly defined mechanisms, MutLγ (comprising MLH1 and MLH3) accumulates at prospective crossover sites and biases dHJ resolution to specifically produce crossovers^17,18^. When MutLγ is dysfunctional, dHJs are still resolved, but with a non-crossover outcome^8^. Consequently, chromosome segregation fails and fertility is diminished. MLH1 and MLH3 are conserved members of the MutL family of DNA mismatch-repair (MMR) factors that couple mismatch-recognition by a MutS complex to downstream excision and resynthesis of the nascent strand^14,16^. MutL complexes from diverse species possess endonuclease activity that provides an initiation site for mismatch excision by a 5’-3 exonuclease such as EXO1^15^. During DNA replication, MutL-catalyzed incision, and thus MMR, is specifically targeted to the nascent strand via an orientation-specific interaction with the replicative clamp, PCNA or the β-clamp in eukaryotes and prokaryotes, respectively^19–22^. Endonuclease activity has been demonstrated for budding yeast and human MutLγ^10–12,23,24^ and the conserved metal-binding active site in MLH3 is required for crossing over in both yeast^7,8^ and mouse^9^. However, how the MutLγ endonuclease is activated and directed to effect crossover-specific resolution of dHJs remains unknown.

### Human MutLγ is an endonuclease

Human MutLγ (hMutLγ) was purified from insect cells (**Fig.1a,b** and **Extended Data Fig. 1**) and endonuclease activity monitored using a supercoiled-plasmid DNA nicking assay (**Fig.1c**). In isolation, hMutLγ displayed a concentration-dependent nicking activity that plateaued at ∼40% of input DNA with 100 nM MutLγ (**Fig.1d–f**). Activity required Mn^2+^ and the metal-binding site of hMLH3, indicating that nuclease activity was specific to hMutLγ. Endonuclease activity of hMutLγ was not seen in Mg^2+^, which is inferred to be the physiological metal cofactor^15,19^ (**Extended Data Fig. 1**). Although Zn^2+^ is present at MutL active sites^15^, nicking was also negligible with Zn^2+^, as well as Ca^2+^, Ni^2+^, Co^2+^ or Cd^2+^ (**Extended Data Fig. 1**). However, these metals were able to compete with Mn^2+^ to inhibit hMutLγ endonuclease activity suggesting that Mn^2+^ is relatively weakly bound, and likely to be non-physiological^15^. The strict dependence on Mn^2+^ is anlogous to human and yeast MutLα complexes (MLH1-PMS2 and Mlh1-Pms1, respectively) when assayed in isolation^19,25^, but contrasts yeast MutLγ, which was reported to show endonuclease activity with a range of metals including Mn^2+^, Co^2+^, Ca^2+^ and Mg^2+ 11^.

**Figure 1.**
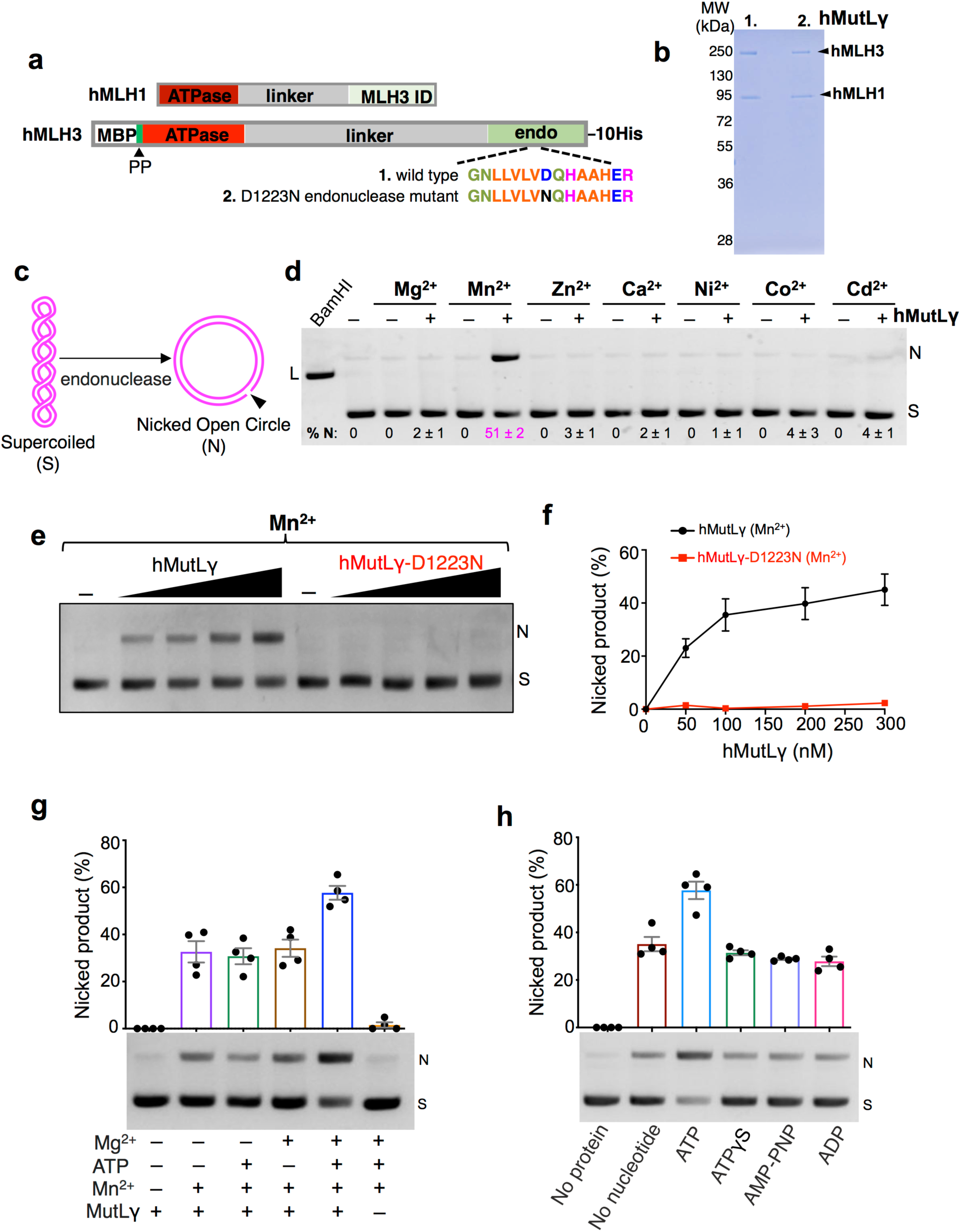
Endonuclease activity of human MutLγ. **a,** Illustration of hMLH1 and hMLH3 expression constructs highlighting domain structure and affinity-purification tags: MLH3 ID, hMLH3 interaction domain; endo, endonuclease domain; MBP, maltose binding protein; PP, PreScission protease cleavage site; 10His, deca-histidine tag. Sequences of the conserved metal binding site for (1) wild-type and (2) the nuclease-dead D1223N mutant of MLH3 are also shown. **b,** SDS-PAGE analysis of purified hMutLγ containing (1) wild-type hMLH3 and (2) mutant hMLH3-D1223N mutant (10% gel stained with Coomassie; also see **Extended Data Fig 1a**). **c,** Supercoiled plasmid nicking assay used to study hMutLγ endonuclease activity. **d,** Representative gel image showing hMutLγ endonuclease assays with various metal ions (100 nM hMutLγ and 5 mM indicated divalent metal ions, incubated at 37°C for 90 min). Migration positions of supercoiled (S) plasmid and nicked (N) product are shown. For reference, plasmid linearized (L) with BamHI is also shown. % N, percent nicked product. Means ± SEM are shown for three experiments after subtracting background nicked product from no-protein controls. **e,** Endonuclease assays for varying concentrations of hMutLγ and hMutLγ-D1223N (0-300 nM protein and 5 mM Mn^2+^ incubated at 37°C for 60 min. **f,** Quantitation of experiments represented in panel **e**. Means ± SEM are shown for three experiments. **g,** Representative gel image and quantification of hMutLγ endonuclease activity with and without ATP and metal cofactors (100 nM hMutLγ, 0.5 mM ATP and 5 mM Mn^2+^ and/or Mg^2+^ incubated at 37°C for 60 min). Means ± SEM are shown for four experiments. **h,** Representative gel image and quantification of hMutLγ endonuclease activity with indicated nucleotides and nucleotide analogs (100 nM hMutLγ, 0.5 mM nucleotide/analog, 5 mM Mn^2+^ and 5 mM Mg^2+^ incubated at 37°C for 60 min). Means ± SEM are shown for four experiments.

MutL proteins are ATPases; binding and hydrolysis of ATP is required for the mismatch correction and meiotic crossover functions of yeast MutLγ *in vivo*^26^, and induces conformational changes *in vitro*^23^. Typically, ATP stimulates MutL endonuclease activity^15^, but surprisingly no stimulation was reported for budding yeast MutLγ, raising the possibility that MutLγ complexes are distinctly regulated^10,11^. On the contrary, we observed that the human MutLγ endonuclease was stimulated 2-fold by ATP, when both Mn^2+^ and Mg^2+^ were present (**Fig 1.g,h** and **Extended Data Fig. 1**). Non-hydrolyzable ATP analogs, ATP-γ-S (adenosine 5′-[γ-thio]-triphosphate) and AMP-PNP (adenylyl-imidodiphosphate), did not stimulate the hMutLγ endonuclease suggesting that ATP hydrolysis is required (**Fig. 1h**).

### hMutSγ stimulates hMutLγ endonuclease activity

The MutSγ complex, MSH4-MSH5, specifically binds JM structures^13,27^ and plays an earlier and more general role than MutLγ in meiotic recombination, localizing to a majority of recombination sites where it stabilizes nascent JMs to facilitate chromosome synapsis^1,17,27,28^. The MMR paradigm predicts that MutSγ may also guide and trigger the MutLγ endonuclease^14,16^. However, MutLγ has its own JM binding activity^10,23^ (below), appears on synapsed chromosomes much later than MutSγ, and localizes only to crossover sites^1,17,28^, raising the possibility that MutLγ acts autonomously during dHJ resolution. Moreover, previous analysis of the yeast enzymes did not detect any stimulation of the MutLγ endonuclease by MutSγ^10^.

By co-immunostaining mouse spermatocyte chromosome spreads for MLH1 and MSH4, we determined that MutLγ closely colocalized with a subset of MutSγ foci as it first emerges at prospective crossover sites in the mid-pachytene stage (**Fig. 2a,b** and **Extended Data Fig. 2**). Co-localization appears to be transient as MSH4 foci subsequently disappeared in late pachytene, while MutLγ persisted. This observation suggests that mammalian MutSγ and MutLγ may transiently interact to modulate MutLγ endonuclease activity. To test this possibility, human MutSγ (hMutSγ) was purified and added to hMutLγ nicking assays (**Fig. 2c–e** and **Extended Data Fig. 3**). In end-point analysis, we observed a modest stimulation (∼20%) of MutLγ endonuclease by MutSγ that was ATP dependent (**Fig. 2d**). Time-course analysis confirmed this inference and revealed an accelerated rate of nicking when MutSγ was present, with product reaching 50% of maximum by 16 min, compared to 22 min for MutLγ alone (**Fig. 2e**). Notably, MutSγ provoked formation of a linearized plasmid product that arose with the same timing as nicked product implying localized concerted incision of both DNA strands by MutLγ.

**Figure 2.**
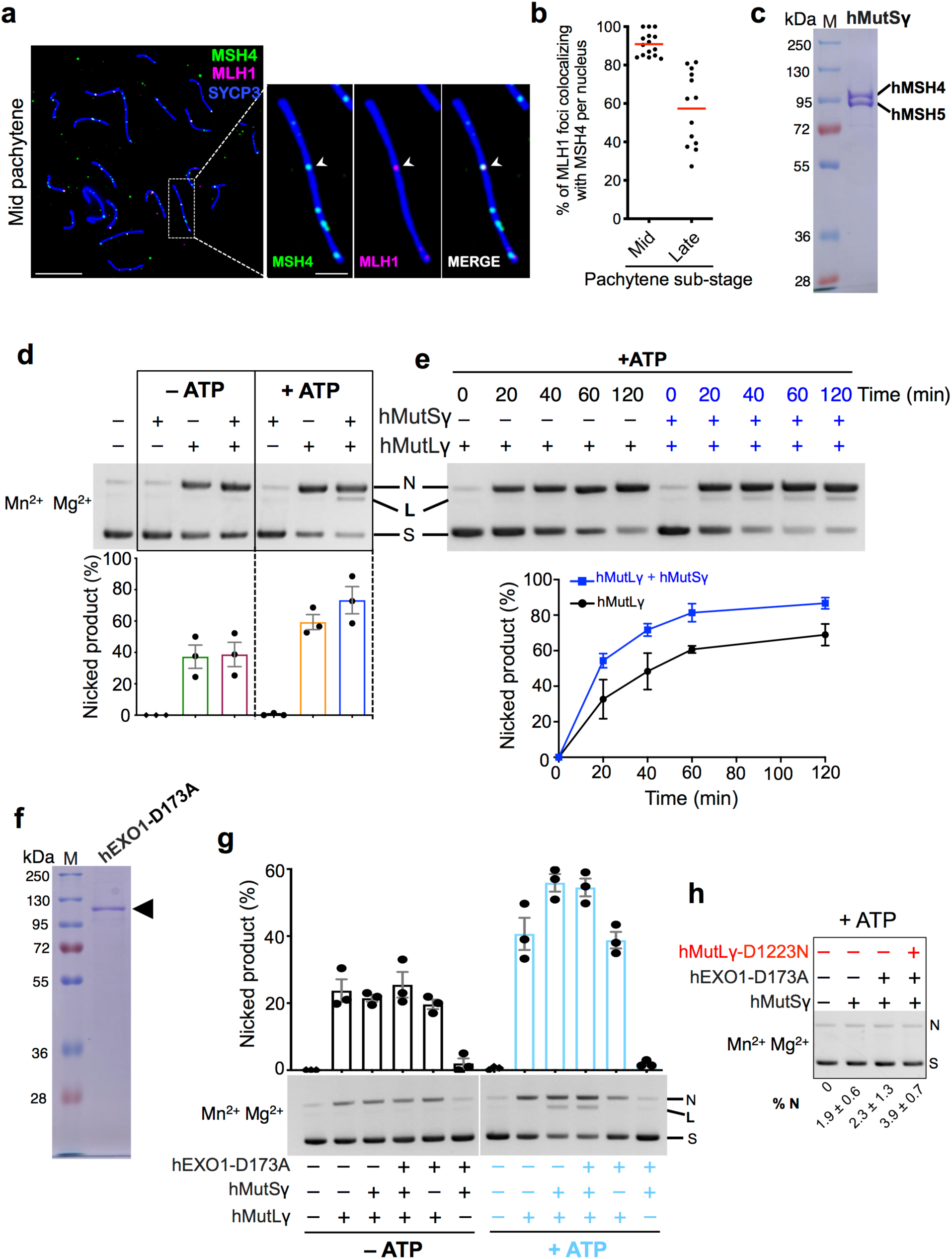
hMutSγ stimulates hMutLγ endonuclease activity. **a,** Representative image of surface-spread chromosomes from a mid-pachytene stage mouse spermatocyte, immuno-stained for MSH4 (green), MLH1 (magenta) and the chromosome axis marker SYCP3 (blue). Magnified panels show an individual pair of synapsed chromosomes in which a single crossover-specific MLH1 focus is colocalized with a MSH4 focus. **b,** Quantification of MLH1-MSH4 colocalization in mid- and late-pachytene stages. **c,** SDS-PAGE analysis of purified hMutSγ (10% gel stained with Coomassie; also see **Extended Data Fig. 3a**). **d,** Representative gel image and quantification of hMutLγ endonuclease activity with and without hMutSγ and ATP (100 nM hMutLγ and hMutSγ, 0.5 mM ATP, 5 mM Mn^2+^ and Mg^2+^, incubated at 37°C for 60 min). Means ± SEM are shown for three experiments. **e,** Time course analysis of hMutLγ endonuclease activity with and without hMutSγ. A representative gel image and quantification of three independent experiments (means ± SEM) are shown. Reaction conditions as described in panel **d**. **f,** SDS-PAGE analysis of purified nuclease-dead hEXO1-D173A (10% gel stained with Coomassie). **g,** Endonuclease assays with varying mixtures of hMutLγ, hMutSγ and hEXO1-D173A, with and without ATP (50 nM each protein, 0.5 mM ATP, 5mM Mn^2+^ and Mg^2+^ incubated at 37°C for 60 min). Means ± SEM are shown for three experiments. **h,** Negative control endonuclease assays with nuclease-dead hMutLγ-D1223N, hMutSγ and hEXO1-D173A Average percent nicking (%N) ± SEM are shown for three experiments.

A nuclease-independent activity of EXO1 facilitates crossing over along the same pathway as MutLγ ^29,30^. We also examined whether nuclease-dead hEXO1-D173A could enhance hMutLγ endonuclease, with or without hMutSγ, but no additional stimulation was seen in the presence of Mn^2+^, Mg^2+^ and ATP (**Fig. 2f–h**, but see below and **Fig. 3**).

**Figure 3.**
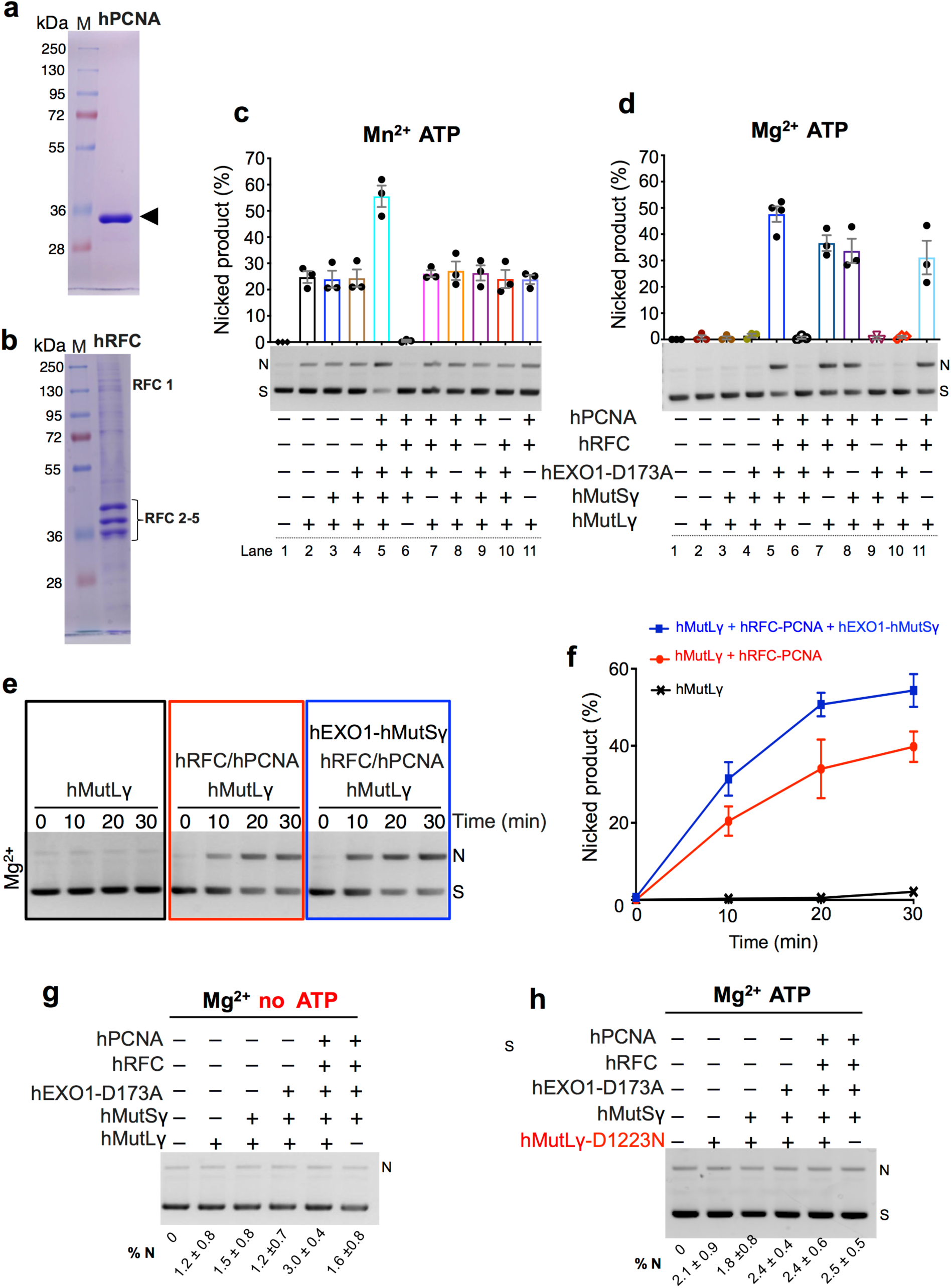
hRFC and hPCNA activate latent hMutLγ endonuclease activity under physiological conditions. **a,b,** SDS-PAGE analysis of purified hPCNA and hRFC (10% gels stained with Coomassie**). c,** Representative gel image and quantification of ensemble endonuclease reactions with Mn^2+^ as the sole metal cofactor. Reactions contained 25 nM indicated proteins, 0.5 mM ATP and 5 mM Mn^2+^ and were incubated at 37°C for 30 min. Means ± SEM are shown for three experiments. **d,** Representative gel image and quantification of ensemble endonuclease reactions with Mg^2+^ as the sole metal cofactor. Reaction conditions were as described in panel **c**, but with 5 mM Mg^2+^ instead of Mn^2+^. **e,f,** Time course analysis of hMutLγ endonuclease activity in Mg^2+^, with and without hRFC/PCNA and hMutSγ-EXO1-D173A. A representative gel image (**e**) and quantification of three independent experiments (**f**, means ± SEM) are shown. Reactions contained 25 nM indicated proteins, 0.5 mM ATP and 5 mM Mg^2+^ and were incubated at 37°C for the indicated times. **g,** No ATP controls for ensemble reactions showing a representative gel image and quantification (means ± SEM). Reaction conditions were as described in panel **d**, but without ATP. **g, h,** Negative control for ensemble endonuclease assays with nuclease-dead hMutLγ-D1223N showing a representative gel image and quantification (means ± SEM). % N, percent nicking.

### RFC-PCNA activates the hMutLγ endonuclease under physiological conditions

Although Mn^2+^ activates MutL endonucleases, its physiological relevance is questionable^15^. In reconstituted mismatch-repair reactions with Mg^2+^ as the sole metal cofactor, MutL complexes are latent endonucleases that are triggered only in the presence of heteroduplex DNA, a cognate MutS complex, and loaded replicative clamp^15,19,21^. When the human clamp, hPCNA, and its DNA loader, hRFC, were added to hMutLγ endonuclease reactions with Mn^2+^ and ATP, nicking was enhanced more than two fold, but only for an ensemble including both hMutSγ and hEXO1-D173A, implying concerted action of all five proteins (**Fig. 3a–c**). Strikingly, hRFC-hPCNA strongly activated the latent hMutLγ endonuclease when Mg^2+^ was the sole metal cofactor (**Fig. 3d**). While hRFC-hPCNA alone could trigger the hMutLγ endonuclease, nicking was further stimulated ∼1.5-fold when both hMutSγ and hEXO1-D173A were also present, i.e. under physiological conditions hMutSγ and hEXO1-D173A appear to act as an interdependent stimulatory factor. Time-course analysis confirmed that MutSγ-EXO1-D173A increased product yield, but not the rate of formation (**Fig. 3e,f**; nicked product reached 50% of maximum by 9 and 10 min, respectively with and without MutSγ-EXO1-D173A). In Mg^2+^-only reactions, activation was absolutely dependent on both hRFC (**Fig. 3d**, lane 9) and ATP (**Fig. 3g**), implying that hPCNA must be loaded onto DNA to activate hMutLγ. Moreover, hPCNA and hMutLγ interacted in solution, suggesting that activation occurs through direct physical interaction between the two proteins (**Extended Data Fig. 4**). Importantly, in these ensemble reactions, all nuclease activity required the metal-binding site of hMLH3, ruling out any contribution from contaminating nucleases (**Fig. 3h**).

### Holliday junctions are specifically bound by hMutLγ and enhance endonuclease activity

Both budding yeast and human MutLγ preferentially bind branched DNA structures, including Holliday junctions (HJs)^10,23^, although Ranjha et al. reported that binding was very sensitive to salt and Mg^2+ 10^. In our hands, hMutLγ selectively bound a broad variety of branched DNA substrates, in the presence of physiological concentrations of salt and Mg^2+^, and excess competitor DNA (poly(dI-dC)) (**Fig. 4a,b**). Four-armed structures bound by hMutLγ included the pro-HJ (pHJ) containing one single-stranded (ss) arm, which approximates an initial D-loop strand-invasion intermediate^13^; a HJ with a 20 nt single-stranded gap in one arm (gHJ), a HJ with a nick at the junction point (nHJ); and a standard HJ (HJ; **Fig. 4a**). HJs and nHJs were bound with very similar affinities (apparent *K_d_* ∼38 and ∼39 nM, respectively; **Fig. 4b**) indicating that additional flexibility around the junction point does not influence hMutLγ binding (unlike, for example, the structure-selective endonuclease Mus81-Mms4^EME1^, ^31^). hMutLγ had significantly higher affinities for the pHJ and gHJ substrates (apparent *K_d_* ∼14 and 16 nM, respectively) suggesting that ssDNA enhances binding (**Fig. 4b**). Indeed, analogous to the budding yeast enzyme ^23^, hMutLγ was able to interact with ssDNA, as shown by its specific binding to a 40 nt poly-dT ssDNA substrate (**Extended Data Fig. 5**). To define the minimal internal ssDNA region that can be efficiently bound by hMutLγ, 80 nt linear DNAs containing single-strand gaps of varying lengths were tested (**Fig. 4c**). Significant binding was not detected for gaps of 5 and 10 nt, but 20 and 40 nt gaps were bound efficiently (**Fig. 4d**). Notably, this minimal binding site is similar to that bound by the canonical ssDNA binding protein RPA^32^. However, RPA binds ssDNA with sub-nanomolar affinity and is therefore predicted to block binding by hMutLγ. Consistently, RPA blocked hMutLγ from binding to a 40 nt gap substrate (**Fig. 4e**). To test whether the branched and ssDNA binding activities are autonomous, gHJ and pHJ substrates were pre-incubated with RPA and then used as binding substrates for hMutLγ (**Fig. 4g**). RPA-gHJ and RPA-pHJ complexes were still readily bound by hMutLγ, but affinities decreased to below that of a HJ (**Fig. 1h**). Thus, as shown by Claeys Bouuaert et al. for yeast MutLγ^23^, the junction and ssDNA binding activities of hMutLγ appear to be largely independent.

**Figure 4.**
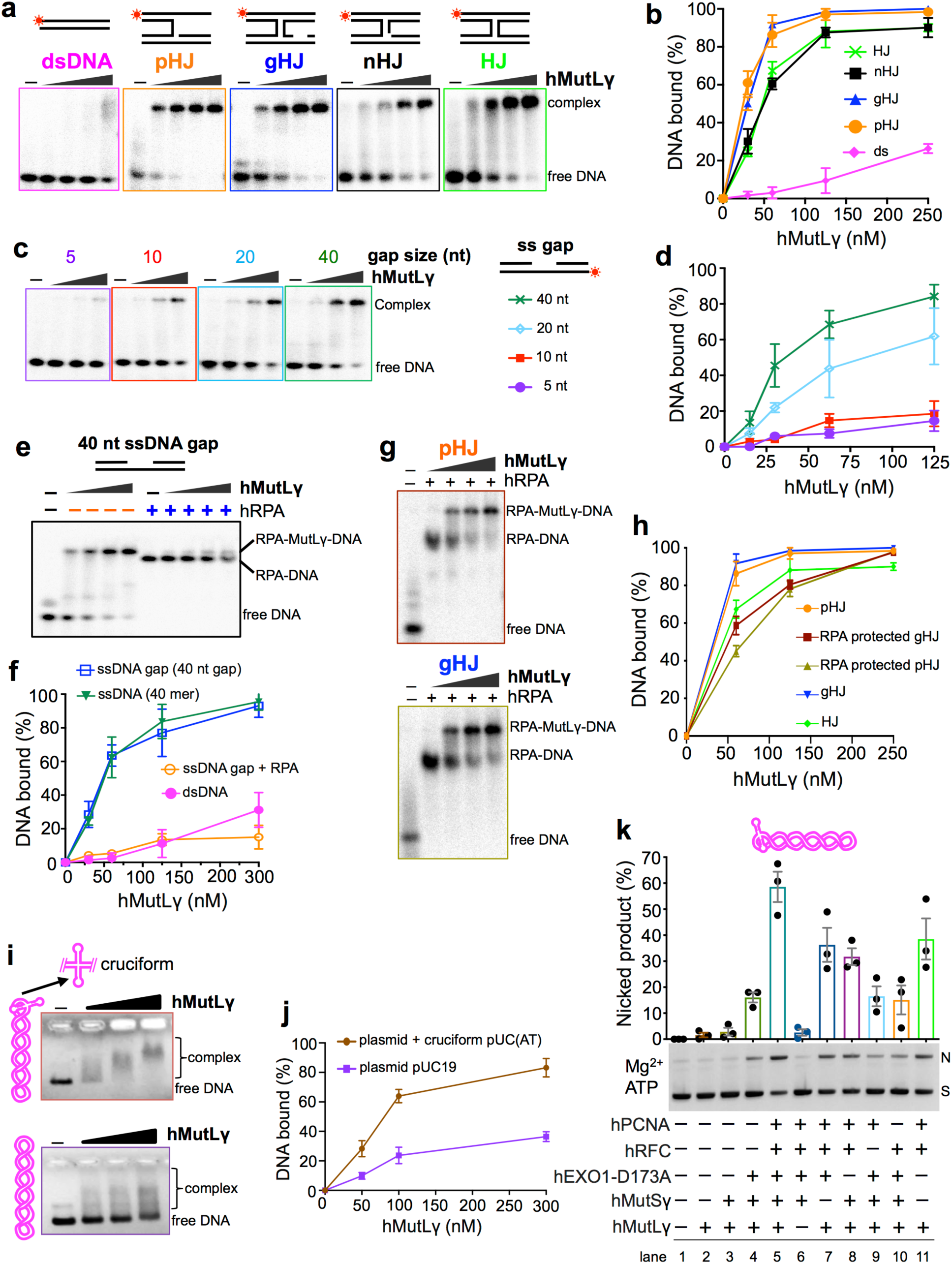
Holliday junctions are specifically bound by hMutLγ and enhance its endonuclease activity. **a,** Representative images of electrophoretic mobility shift assays (EMSAs) analyzing hMutLγ binding to control double-stranded DNA and the illustrated branched DNA structures. Terminally ^32^P-labeled strands are indicated by asterisks. **b,** Quantification of the EMSAs represented in panel **a** showing means ± SEM for three independent experiments. **c,** Representative images of EMSAs analyzing hMutLγ binding to 80mer DNAs containing single-strand gaps of varying size. **d,** Quantification of the EMSAs represented in panel **c** showing means ± SEM for three independent experiments. **e,** Representative images of EMSAs for hMutLγ binding to an 80mer DNA containing a 40 nucleotide gap, with or without prior incubation with RPA. **f,** Quantification of the EMSAs represented in panel **e** and **Extended Data Fig. 5** shows that hMutLγ binds a 40 nt gap with almost identical affinity as a 40mer single stranded DNA; and that pre-incubation with RPA blocks hMutLγ binding. Error bars show means ± SEM for three independent experiments. **g,** Representative images of EMSAs for hMutLγ binding to pro-HJs and gapped-HJs that were pre-incubated with RPA. **h,** Quantification of the EMSAs represented in panels **a** and **g**. Error bars show means ± SEM for three independent experiments. **i,** Representative images of EMSAs for hMutLγ binding to pUC(AT) and pUC19 supercoiled plasmid substrates. **j,** Quantification of the EMSAs represented in panel **i**. Error bars show means ± SEM for three independent experiments. **k,** Representative gel image and quantification of ensemble endonuclease reactions with using pUC(AT) as substrate and Mg^2+^ as the sole metal cofactor. Reactions contained 25 nM indicated proteins, 0.5 mM ATP and 5 mM Mg^2+^ and were incubated at 37°C for 30 min. Means ± SEM are shown for three experiments.

Despite the intrinsic JM binding ability of hMutLγ, resolution or even nicking of model HJ-containing substrates was not detected (**Extended Data Fig. 6**) implying that hMutLγ cannot catalyze symmetric incisions across the junction, i.e. is not a classical HJ resolving enzyme^33^. However, the presence of a HJ did influence the binding and incision of a plasmid substrate (**Fig. 4i–k** and **Extended Data Fig. 7**). pUC(AT) contains an (AT)_20_ inverted repeat that extrudes into a four-way hairpin junction upon supercoiling, but is otherwise identical to pUC19^34^. By agarose-gel electrophoresis, a heterogeneous smear of hMutLγ–pUC19 complexes was observed suggesting unstable binding (**Fig. 4i**). By contrast, hMutLγ–pUC(AT) complexes migrated as relatively discrete species and overall efficiency of stable complex formation was much higher (**Fig. 4j**). Moreover, the decreasing mobility of hMutLγ–pUC(AT) complexes observed with increasing hMutLγ concentration points to formation of higher-order complexes analogous to the cooperative, multimeric oligomers described for yeast MutLγ^12,23^.

In ensemble plasmid-nicking reactions with Mg^2+^ and ATP, pUC(AT) was incised up to ∼23% more efficiently than pUC19 (**Fig. 4k**, lanes 5 and 11; and **Extended Data Fig. 7**). Analogous to pUC19, pUC(AT) cleavage was most efficient when hMutLγ, hMutSγ, hEXO1-D173A and hRFC-hPCNA were present, with hMutSγ and EXO1-D173A appearing to function interdependently. Distinct from pUC19, however, pUC(AT) incision was not absolutely dependent on hRFC-hPCNA, with ∼15% nicked product being formed in reactions containing hMutLγ, hMutSγ and hEXO1-D173A (**Fig. 4k**, lanes 4, 9 and 10). Thus, the HJ in pUC(AT) modulates both the binding and incision of DNA by hMutLγ.

### RFC-PCNA facilitates MutLγ-dependent crossing over

A role for RFC-PCNA in MutLγ−dependent crossing over was demonstrated *in vivo* in budding yeast (**Fig. 5**). An auxin-inducible degron (AID) allele of Rfc1 was employed to inactivate RFC, and thereby prevent PCNA loading, precisely at the time of dHJ resolution (**Fig. 5a**; note that AID alleles of PCNA were not functional). An inducible allele of the *NDT80* gene allowed reversible arrest of cells at the pachytene stage of meiosis, in which chromosomes are fully-synapsed and dHJs are poised for resolution^35,36^. Following release from arrest and addition of auxin, Rfc1-AID was rapidly degraded (**Fig. 5b,c**) and crossing over at a recombination hotspot, monitored by Southern analysis, was reduced from 17.5% (± 2.2% SEM) to 10.4% (± 1.8% SEM) (**Fig. 5d,e**). Similar analysis of *RFC1-AID mlh3Δ* cells indicated that RFC promotes primarily MutLγ−dependent crossovers. However, approximately one-third of *MLH3*-independent crossovers also appeared to be RFC dependent, which may be explained by the ability of RFC-PCNA to also stimulate the structure-selective endonuclease, Mus81-Mms4^EME1 37^, which defines a second crossover pathway in budding yeast^8^. Rfc1-AID degradation did not alter the timing or efficiency of dHJ resolution indicating that the crossover outcome, but not resolution *per se*, is facilitated by RFC-PCNA (**Fig. 5f**, and **Extended Data Fig. 8**). Under these conditions, *mlh3Δ* mutation did delay dHJ turnover, but resolution was ultimately efficient and Rfc1-AID degradation did not further alter the timing or efficiency of resolution.

**Figure 5.**
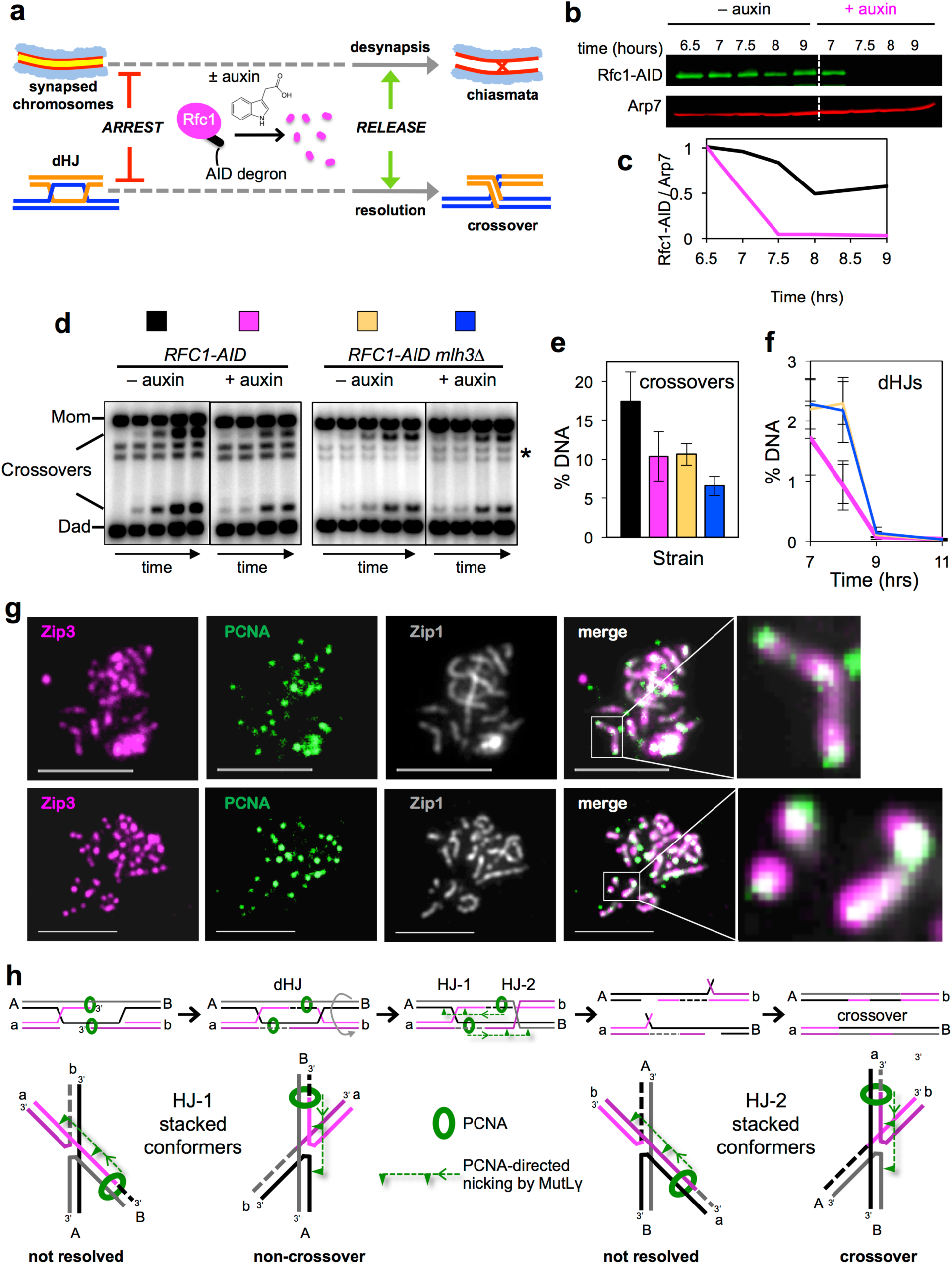
RFC-PCNA promotes crossing over *in vivo*. **a,** Regimen to degrade Rfc1-AID during the late stages of meiotic prophase. **b,** Western image showing Rfc1-AID levels in cells with and without the addition of auxin. **c,** Quantification of Rfc1-AID protein levels relative to ab Arp7 loading control. **d,** Representative Southern blot images of crossover analysis at the *HIS4::LEU2* recombination hotspot. Time points are 0, 7, 8, 9, and 11 hours after induction of meiosis. **e,f,** Quantification of crossovers products and double Holliday junction (dHJ) intermediates. “% DNA” is percentage of total hybridizing DNA signal on 1D Southern blots, with error bars representing means ± SEM. **g,** Representative images of surface spread meiotic nuclei immunostained for PCNA (green), the crossover marker Zip3 (magenta), and synaptonemal complex protein Zip1 (grey). Scale bars = 5 μm. PCNA and Zip3 foci were quantified in nuclei with pachytene morphologies (fully synapsed chromosomes indicated by linear Zip1 staining). Magnified panels show individual synaptonemal complexes with colocalizing PCNA and Zip3 signals. **h,** Model of crossover-specific dHJ resolution during meiosis. Top row: the orientations of PCNA molecules loaded during dHJ maturation direct MutLγ to catalyze strand-specific nicking on either side of the two HJs. This pattern of incisions, in combination with migration about the junctions, results in crossover-specific resolution. Bottom row: PCNA-directed MutLγ incision results in resolution for only one of the two stacked-X HJ conformers, which may be favored by factors such as MutSγ. Alternatively, conformers may freely interconvert. For HJ-1, resolution always produces a non-crossover; while for HJ-2, crossover products are always formed. See text for further details.

Consistent with a role for RFC-PCNA in crossing over, PCNA localized to a subset of prospective crossover sites marked by the Zip3 protein (**Fig. 5g**)^38^. In chromosome spreads from cells arrested in pachytene, PCNA and Zip3 immunostaining foci averaged 15.4 (± 1.1 SEM) and 47.2 (± 1.3 SEM), respectively (n=25 nuclei); and 85.1% (± 1.9% SD) of PCNA foci colocalized with Zip3.

## Discussion

Asymmetric incision is required for the crossover-biased resolution of dHJs during meiosis, but how asymmetry is imposed has remained unclear. The requirements for MutSγ, MutLγ and EXO1 in meiotic crossing over evoked a MMR-like mechanism, involving MutLγ-catalyzed nicking of dHJs^1,17,18^. Our discoveries implicating RFC-PCNA in MutLγ-dependent crossing over strongly suggest that dHJ resolution is indeed fundamentally analogous to canonical MMR. We propose a specific model for crossover-specific dHJ resolution in which orientation-specific loading of PCNA, during the DNA synthesis associated with dHJ formation, directs MutLγ to incise specific DNA strands on both sides of the junction points (**Fig. 5h**). In reconstituted MMR reactions, PCNA, loaded at a nick, signals over hundreds of base pairs to direct incision of the same non-continuous strand by MutLα^14–16^. While incisions are biased upstream of the mismatch, in order to initiate excision by the 5’-3’ exonuclease EXO1, nicking also occurs at downstream locations. By analogy, we propose that the PCNA molecules involved in recombination-associated DNA synthesis direct strand-specific nicking by MutLγ on both sides of the two HJs (**Fig. 5h**). Resolution occurs only when the HJs adopt one of the two co-axially stacked-X conformers, which could be favored by the binding of MutSγ^27^ or other pro-crossover factors, and always specifies a crossover outcome. We suggest that the near continuous duplexes formed by the stacked HJ arms enable the strand-specificity of PCNA-directed MutLγ-catalyzed incision to be maintained across the junction point, even though the targeted strands are not contiguous. Given that nicking occurs some distance from the exchange points, our model predicts that junction migration via a helicase will be required for crossover-specific dHJ resolution, consistent with the known role of the RecQ-family helicase Sgs1/BLM^8,39^. Our model provides a solution to the critical question of how a crossover outcome is enforced at designated sites to help ensure accurate chromosome segregation during meiosis.

## Methods

### Purification hMutLγ

MBP-hMLH3-His ^10^ and untagged hMLH1 ^19^ were co-expressed in insect cells according to standard protocols (Bac-to-Bac system, Invitrogen) and hMutLγ was purified as described by Ranjha et al.^10^ with the following modifications. *Spodoptera frugiperda* Sf9 cells were coinfected with optimized ratios of baculoviruses, cells were harvested 52 h after infection, washed with phosphate-buffered saline (PBS), frozen in liquid nitrogen and stored at −80°C. A typical purification was performed with cell pellets from 3.6 L of culture. All subsequent steps were carried out at <4°C. Cells were resuspended in 3 volumes of lysis buffer (50 mM HEPES-NaOH pH 7.4, 1 mM DTT, 1 mM EDTA, 1:500 (v/v) protease inhibitor cocktail (P8340 Sigma), 1 mM phenylmethylsulfonyl fluoride, 30 µg/ml leupeptin) and stirred slowly for 15 min. Glycerol was added to 16% and 5 M NaCl was added for an final concentration of 325 mM. The sample was stirred for 30 min and then centrifuged at 50,000 g for 30 min. The clarified extract was bound in batch mode to 8 ml of pre-equilibrated amylose resin (New England Biolabs) for 1 hr. The resin was then washed extensively with 300 ml of buffer a buffer comprising 50 mM HEPES-NaOH pH 7.4, 2 mM β-mercaptoethanol, 250 mM NaCl, 10 % glycerol, 1 mM phenylmethylsulfonyl fluoride, 10 µg/ml leupeptin. Bound protein was eluted in wash buffer containing 10 mM maltose. Next, the MBP tag was cleaved from MLH3 with PreScission protease at a 1:16 ratio of proteins for 1 hr. The sample was applied to a pre-equilibrated 1 ml nickel-nitrilotriacetic acid resin column (Ni-NTA, Qiagen) over a 45 min period in wash buffer supplemented with 20 mM imidazole. The column was then washed under gravity flow with 100 ml buffer containing 60 mM imidazole, before eluting bound protein with the same wash buffer containing 400 mM imidazole. Pooled fractions were dialyzed against a buffer containing 25 mM HEPES-NaOH pH 7.4, 1 mM DTT 150 mM NaCl, 20% glycerol, 0.5 mM phenylmethylsulfonyl, concentrated with 50 kDa cut-off Amicon centrifugal filters (Millipore), and stored at 4 or 80 °C. Nuclease deficient hMutLγ (containing hMLH3-D1223N) was created by using a QuikChange II Site-Directed Mutagenesis Kit (Agilent Technologies Inc, 200524). hMutLγ containing hMLH3-D1223N protein was prepared in the same way as wild-type protein. Protein concentrations were determined by Bradford assay using Bovine serum albumin as a standard, and by spectroscopic absorption at 280 nm.

### Purification of hMutSγ

*hMSH4* and *hMSH5-6His* coding regions were synthesized (GeneCopoeia), digested with SmaI and KpnI (for *MSH4*) or SalI and XbaI (for *MSH5-6His*) and cloned into the pFastBac Dual vector (ThermoFisher). Baculoviruses were prepared according to manufacturer’s instructions and hMutSγ was purified as described previously^40^. Insect cells were infected with hMSH4/hMSH5-6His baculovirus at 1×10^6^ per mL, and incubated at 27°C for 56 hrs with shaking. Cells were centrifuged at 2000 rpm for 10 minutes at 4°C, washed with ice cold PBS, frozen in liquid nitrogen and stored at −80°C. A typical purification was performed with cell pellets from 1.6 L of culture. All subsequent steps were carried out at <4°C. Cells were thawed on ice-cold water for 5 min and then resuspended in 4 volumes of lysis buffer (25 mM HEPES pH 8.1, 300 mM NaCl, 20 mM imidazole, 10% glycerol, 500 uM PMSF, 10 µg/mL Pepstatin (Sigma), 10 µg/mL Leupeptin (Sigma), 10 µg/mL Aprotinin (Sigma), 1 µg/ml E-64 (Sigma) and lysate was passed five times through a 25G needle. The lysate was centrifuged at 35,000 rpm for 1 hour at 4°C, and clarified extract was loaded onto a 5 ml Ni-NTA column (GE Healthcare) pre-equilibrated with Buffer A (25 mM HEPES-NaOH pH 8.1, 300 mM NaCl, 20 mM Imidazole, 10% glycerol containing 500 µM PMSF, 1 µg/mL Pepstatin, and 1 µg/mL Leupeptin). The column was washed with 20 column volumes of Buffer A and then protein was eluted with a linear gradient of imidazole (20-300 mM) in Buffer A. Peak fraction were pooled and applied to PBE-94 (Pharmacia) and Heparin Sepharose (GE Healthcare) columns in series. PBE/Heparin flow through was dialyzed against Buffer B (HEPES-NaOH pH 7.8, 100 mM NaCl, 1 mM DTT, 0.1 mM EDTA, and 10% glycerol, containing 500 µM PMSF, 1 µg/mL Pepstatin, and 1 µg/mL Leupeptin). The protein was then loaded onto a 1 ml Heparin-Sepharose column (GE Healthcare) pre-equilibrated with Buffer B and eluted with a linear gradient of NaCl (100 mM to 1 M). Peak fractions containing hMutSγ were pooled and dialyzed against Buffer C (25 mM HEPES-NaOH pH 7.8, 100 mM NaCl, 1 mM DTT, 0.1 mM EDTA, 20% glycerol) and concentrated with 50 kDa cut-off Amicon centrifugal filters (Millipore). Aliquots were frozen in liquid nitrogen and stored at −80°C. Protein concentation was determined as described above.

### Purification of hEXO1-D173A

hEXO1 (D173A) was prepared as described previously^19,40^. Sf9 insect cells were infected with baculovirus expressing hEXO1-D173A at 1 x 10^6^ per mL and incubated at 27°C for 58 hrs. Cells were centrifuged at 2000 rpm for 10 minutes at 4°C, frozen in liquid nitrogen and stored at −80°C. All subsequent steps were carried out at <4°C. Cells were thawed on ice-cold water for 5 min and then resuspended in 5 volumes of lysis buffer A (20 mM KPO_4_ pH 7.5, 150 mM KCL, 0.1 mM EDTA) with Protease inhibitors (500 uM PMSF, 1 ug/mL Pepstatin, 10 ug/mL Leupeptin, 10 ug/mL Aprotinin, 1ug/ml E-64). The cells were lysed using a dounce homogenizer for 5 strokes followed by gentle mixing on ice and addition of 50 % glycerol to a final concertation 10%, and 2 M KCl for a final concentration 150 mM. The suspension was then incubated for an additional 15 min with gentle stirring. The extract was then clarified by centrifuged at 55,000 rpm for 30 min and loaded onto a 15-ml Q-Sepharose column (GE Healthcare), preequilibrated with Q-Sepharose buffer (20 mM KPO_4_, pH 7.5, 100 mM KCl, 0.1 mM EDTA, 10% glycerol containing the same protease inhibitors used in the lysis buffer). The column was washed with 10 column volumes of Q-Sepharose buffer and eluted with a 5-column gradient of KCI gradient (100-400 mM) supplemented with 1mM DTT. hEXO1-D173A peak fractions were pooled and diluted two-fold with buffer B (20 mM KPO_4_ pH 7.5, 150 mM KCL,10% glycerol 0.1 mM EDTA,1 mM DTT 0.5 mM PMSF, 1 ug/mL Pepstatin, 1ug/mL Leupeptin, 1 ug/mL Aprotinin) and fractionated on a 5 ml Heparin column (GE Healthcare) pre-equilibrated with buffer B. The column was washed with 20 column volumes of buffer B and eluted with a 10-column gradient of KCl (100-415 mM). Peak fraction were diluted to 150 mM KCl with buffer B and loaded onto a 1 ml MonoS column (GE Healthcare) pre-quilibrated with Mono S Buffer (20 mM KPO_4_, pH 7.5, 150 mM KCL, 0.1 mM EDTA 10% glycerol, 0.5 mM PMSF, 1mM DTT, 1 ug/mL Pepstatin, 1 ug/mL Leupeptin, 1 ug/mL Aprotinin). Non-specifically bound protein was removed with Mono S Buffer and elution was performed with a 2-column volume linear KCl gradient (100-500 mM). hEXO1-D173A eluted at ∼300 mM KCl on both the Heparin and MonoS columns. Peak fractions were subjected to Gel filtration chromatography on a S200 column (GE Healthcare) using S200 Buffer (20 mM KPO_4_, pH 7.5, 300 mM KCL, 0.1 mM EDTA, 5% Glycerol, 0.5 mM PMSF and 1 ug/mL Leupeptin). Peak fractions were collected, supplemented with 20% glycerol and 1 mM DTT, flash-frozen in liquid nitrogen, and stored at −80  °C.

### hPCNA and hRFC and hRPA

Purified hPCNA, hRFC and hRPA were generous gifts of Paul Modrich (Duke University), Jerry Hurwitz (MSKCC), and Steve Kowalczykowski (UC Davis), respectively.

### DNA substrates

All oligonucleotides were obtained from Integrated DNA Technologies. Sequences are listed in Table 1. Oligonucleotides were 5’ end-labeled with [γ-32P] ATP (PerkinElmer Life Sciences) using T4 polynucleotide kinase (New England Biolabs). Unincorporated nucleotide was removed using G-50 Micro Columns (GE Healthcare). Substrate DNAs were annealed by combining end-labeled oligonucleotide with a 2-fold molar excess of the unlabeled complementary oligos. The end-labeled DNA substrates were then gel purified. Negatively supercoiled pUC19 and pUC-AT plasmids (New England Biolabs) were prepared using standard protocols.

### Nicking endonuclease assays

Endonuclease nicking assays were performed using 100 ng negatively supercoiled pUC19 or cruciform-containing pUC-AT plasmid DNA in 20 µl reaction mixtures containing 25 mM HEPES-NaOH pH 7.4, 50 mM NaCl, 1mM DTT, 5% glycerol, 0.2 mg/mL bovine serum albumin (NEB), indicated concentrations of purified proteins, and 5 mM metal cofactors (MnCl_2_, MgCl_2_, ZnCl_2_, CaCl_2_, NiCl_2_, or CoCl_2_, SIGMA). Where indicated, 5mM ATP, ADP, AMP-PNP or ATP-γ-S (Sigma) was added.

Reactions employing RFC-PCNA contained 25 mM HEPES-NaOH pH 7.4, 25 mM NaCl, 1mM DTT, 0.2 mg/mL bovine serum albumin BSA, 2% glycerol and 5 mM Mg^2+^ or Mn^2+^. 25 nM hRFC, hPCNA, hEXO1-D173A, hMutLγ and hMutSγ and 0.5 mM ATP were added as indicated. Reactions were assembled on ice, incubated at 37°C for indicated times, and stopped and deproteinized by adding 4 µl Stop buffer (200 mM EDTA and 2 mg/ml proteinase K, 2% SDS) and incubating for 20 min at 55°C). Plasmid isoforms were separated by electrophoresis in 1% agarose gel on 1% TAE (1.25 V/cm ∼ 2 hrs) and gels were stained with 0.5 µg/ml ethidium bromide. Gel images were acquired with a Fluro Chem 8900 (Alpha Innotech) and quantified using Image J.

### EMSA DNA binding assay

Indicated concentrations of hMutLγ and 1 nM of ^32^P-labeled DNA substrate were incubated in a 10 μl reaction mixture containing 25 mM HEPES (pH 7.8), 100 mM NaCl, 5 mM MgCl_2,_ 5% glycerol, 0.1 mg/ml BSA, 2 ng/µl poly(dI/dC) (SIGMA) and 1 mM DTT supplemented with indicated concentrations of nucleotides and/or divalent metal ions. Reaction mixtures were assembled on ice and incubated at 4°C for 10 min. To assemble hRPA coated ssDNA substrates, 1 nM of ^32^P-labeled DNA substrates were combined in the reaction buffer described above together with indicated concentrations of hRPA and pre-incubated for 10 min at 4°C before adding hMutLγ and incubating for an additional 10 min. Bound and free DNA species were resolved by electrophoresis using non-denaturing 5% polyacrylamide gels in TAE buffer. Gels were dried on DE81 paper (Whatman) exposed to storage phosphor-storage screens and imaged using a Typhoon FLA 9000 imager (GE Healthcare). Fractions of mobility-shifted substrate were quantified, plotted and analyzed using Image J and GraphPad Prism software.

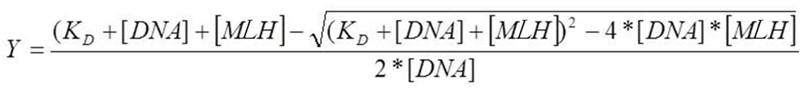

The apparent equilibrium binding constants were obtained by fitting the data to the 1:1 binding equation, above, where *K*_D_ is the equilibrium dissociation constant, [DNA] is the concentration of DNA substrate (1 nM) and [MLH] is concentration of hMutLγ.

### Denaturing urea PAGE analysis of HJ endonuclease reactions

Nuclease assays with 1 nM labeled HJ or nHJ substrates were performed in 15 μl volumes in presence of 5 mM Mg^2+^ and/or Mn^2+^, 0.5 mM ATP, and 25 nM each protein in nuclease reaction buffer (above) for 30 min at 37°C. Reactions were stopped by the addition of EDTA (50 mM) and Proteinase K treatment was performed for 15 min at 55°C. 15 μl formamide dye (98% formamide, 10 mM EDTA, 0.1% bromophenol blue and 0.1% and xylene cyanol) was added and reactions were denatured at 95°C for 5 min. The denatured products were separated on 15% denaturing polyacrylamide gel (National diagnostic lab Urea PAGE system) in 1X TBE buffer. The gels were rinsed twice in a solution containing 40% methanol, 10% acetic acid, and 5% glycerol, and then fixed in the same solution for 30 min at room temperature. Gels were then vacuum dried on 3 mm Chr Whatman paper, exposed to storage phosphor screen and scanned using a Typhoon Phosphor imager (FLA 9000, GE Healthcare).

### Analysis of hMutLγ–hPCNA interaction

0.6 µg anti-PCNA antibody (Santa Cruz Biotechnology F2, 25280) was incubated with 20 µl Protein A beads and equilibrated in PBS buffer at 4°C for 2 hr with gentle mixing. The beads were washed twice with 100 µl PBS, mixed with 1 µg recombinant hPCNA and 0.5 µg hMutLγ in 100 µl binding buffer (20 mM HEPES pH7.4, 50 mM NaCl, 5% glycerol, 0.1 mM EDTA, 1 mM DTT), and incubated overnight at 4°C with gentle mixing. Supernatant was collected as unbound protein following a gentle spin at 1000 rpm for 2 min. Beads were washed three times with binding buffer (20 mM HEPES 7.4, 50 mM NaCl, 5% Glycerol, 0.1 mM EDTA, 1 mM DTT). The bound protein complexes were eluted in Laemmli buffer for 3 min at 95°C and separated on a 10% SDS-PAGE gel. The gel was rinsed with water, stained with SYPRO-Ruby (Lonza-50562) according to manufacturers recommendations, and imaged using a Typhoon FLA 9000 imager (GE Healthcare).

### Mice

Mice were congenic with the C57BL/6J background and maintained and used for experiments according to the guidelines of Institutional Animal Care and Use Committees of the University of California Davis.

### Mouse cytology

Surface-spread chromosomes of mouse spermatocytes were prepared from 18 days postpartum male mice using the drying-down technique as previously described^41^. Immuno-staining was performed as previously described^41^ using the following primary and secondary antibodies: goat anti-SYCP3 (Santa Cruz Biotechnology, sc-20845; 1:200), rabbit anti-MSH4 (a gift from Dr. Paula Cohen; 1:200), mouse monoclonal anti-MLH1 (4C9C7) (Cell Signaling Technology, 3515S; 1:30), AMCA anti-goat (Jackson ImmunoResearch, 705-155-147; 1:50), Cy3 anti-rabbit (Jackson ImmunoResearch, 711-165-152; 1:100), and AlexaFluor647 anti-mouse (Jackson ImmunoResearch, 715-605-151; 1:100).

### Image acquisition and analysis of mouse cytology

Chromosome spreads were imaged using a Zeiss Axioplan II microscope equipped with a 63 x Plan-Apochromat 1.4 NA objective, EXFO X-Cite metal halide light source, and a Hamamatsu ORCA-ER CCD camera. Images were processed and analyzed using Volocity software (Perkin Elmer). Focus counts and colocalization were determined manually. Early, mid and late pachytene stages were defined using standard criteria.

### Yeast strains

Full genotypes of the *Saccharomyces cerevisiae* strains (SK1 background) used in this study are listed in **Supplementary Table 2**. Adaptation of the Auxin-Induced Degron (AID) system for use during meiosis has been described previously^42,43^. Fusion of a minimal *AID* cassette to *RFC1* was constructed using plasmid pHyg-AID*-9Myc as a template for PCR-mediated allele replacement, as previously described^36,44^. The estrogen-inducible *IN-NDT80 GAL4-ER* system has been described^35,36,45,46^.

### Meiotic time courses and DNA physical assays

Detailed protocols for meiotic time courses and DNA physical assays at the *HIS4::LEU2* locus have been described^47,48^. Degradation of Rfc1-AID was performed as pachytene cells were released from *IN-NDT80* arrest as described previously^43^, with the following modifications: at 6.5 hours after induction of meiosis, CuSO_4_ (100 mM stock in dH_2_O) was added for a final concentration of 50 μM to induce expression of *P_CUP1_-OsTIR1*. Then at 7 hours, auxin (3-indoleacetic acid, Sigma 13750, 2 M stock in DMSO) was added to one subculture at a final concentration of 2 mM to induce degradation of Rfc1-AID; and estradiol (5 mM stock, Sigma E2758 in ethanol) was added to both subcultures to induce *IN-NDT80*. At 7.5 hours, auxin was added again at 1 mM. Cell samples were collected to assay protein depletion, meiotic divisions, and recombination intermediates as described previously^36^.

### Western analysis

Whole cell extracts were prepared using the TCA extraction method, essentially as described^43^. Samples were analyzed by standard SDS-PAGE and Western blotting. Anti-c-myc monoclonal antibody (Roche; 11667149001) was diluted 1:1,000 to detect Rfc1-AID; Arp7 polyclonal antibody (Santa Cruz, SC-8961) was used at 1:5,000 dilution as a loading control. Secondary antibodies, used at 1:5,000 dilution, were IRDye® 800CW Donkey anti-Mouse IgG (LI-COR, 925-32212) and IRDye® 680LT Donkey anti-Goat IgG (LI-COR, 925-68024). Membranes were imaged with an Odyssey Infrared Imager (LI-COR). Quantification of protein bands was performed using Image Studio Lite 5.0.21 software.

### Yeast cytology and chromosome spreading

Meiotic cells were collected at indicated time points and fixation and chromosome spread was performed as described using 4% paraformaldeyde^49^. Immunostaining was performed as described^49^. Primary antibodies were anti-PCNA (Abcam; ab70472, 1:100), anti-Zip3 antibody (a gift from Dr. Akira Shinohara, 1:400) and anti-Zip1 (a gift from Dr. Scott Keeney, 1:400); all incubated overnight at room temperature in 100 ul TBS/BSA buffer (10 mM Tris PH7.5, 150 mM NaCl, 1% BSA) Secondary antibodies were anti-rabbit 568 (A11036 Molecular Probes, 1:1000), anti-mouse 488 (A11029 Molecular Probes, 1:1000), anti-rabbit 647 (A21245 Invitrogen), and anti-guinea pig 555 (A21435 Life Technologies); all for 1 hr at 37°C. Coverslips were mounted with Prolong Gold antifade reagent (Invitrogen, P36930). Digital images were captured using a Zeiss Airyscan LSM800 with Axiocam and analyzed using Zen (blue edition); or a Zeiss Axioplan II microscope, Hamamatsu ORCA-ER CCD camera and analyzed using Volocity software. Co-localization of protein foci was assigned to overlapping foci. Scatterplots were generated using the GraphPad program in Prism.

## Acknowledgements

We thank Paul Modrich (Duke), Jerry Hurwitz (MSKCC), Ken Marians (MSKCC), Steve Kowalczykowski (UCD) Paula Cohen (Cornell), Scott Keeney (MSKCC) and Akira Shinohara (Osaka University) for reagents; Petr Cejka (Università della Svizzera italiana) for reagents and communicating unpublished data; and members of the Hunter Lab for support and discussions. This work was supported by NIH NIGMS grant GM074223 to N.H. S.O. was supported by NIH NIGMS T32 Training Program in Molecular and Cellular Biology 5T32GM007377 and an F31 Ruth L. Kirschstein National Research Service Award 1F31GM125106. M.I. was supported by a Japan Society for the Promotion of Science postdoctoral fellowship for research abroad. N.H. is an Investigator of the Howard Hughes Medical Institute.

## Author Contributions

D.S.K., S.O., M.H. and N.H. conceived the study and designed most experiments. All authors performed experiments and analyzed the data. D.S.K., S.O and N.H. wrote the manuscript with inputs and edits from all authors.

## Supplementary information

This file contains Supplementary Tables 1 and 2.

## Extended Data

**Extended Data Figure 1.**
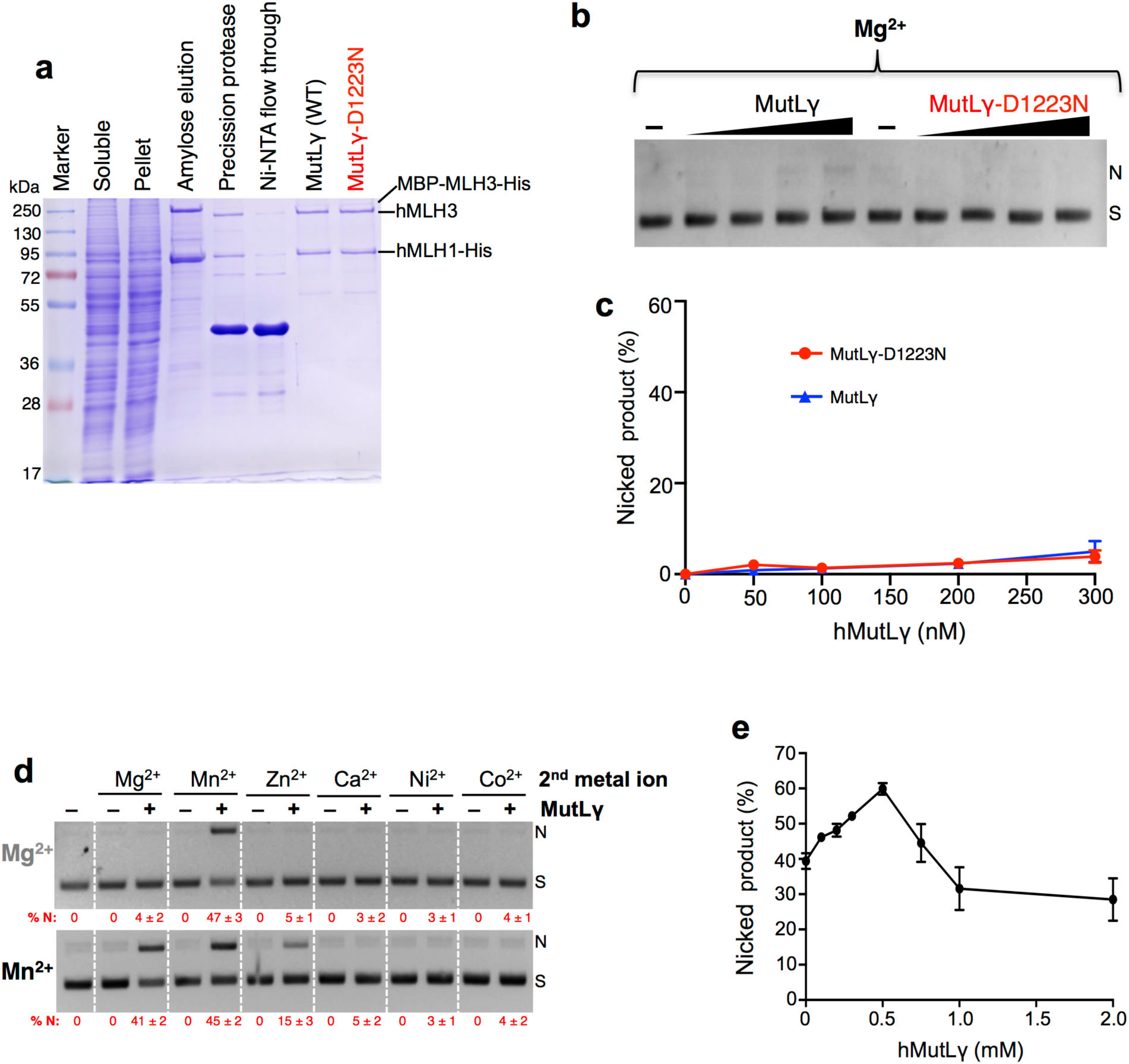
Purification of human MutLγ and characterization of endonuclease experiment. **a,** Representative MutLγ purification steps monitored by 10% SDS-PAGE stained with Coomassie blue. Amylose enriched fractions were treated with Precission Protease, to cleave the maltose-binding protein tag, and then subjected to Ni-NTA affinity purification. **b,** Representative endonuclease assays for varying concentrations of hMutLγ and hMutLγ-D1223N (0-300 nM protein and 5 mM Mg^2+^ incubated at 37°C for 60 min). Migration positions of supercoiled (S) plasmid and nicked (N) product are indicated. **c,** Quantitation of experiments represented in panel **b** shows that the hMutLγ endonuclease is inactive in Mg^2+^. Means ± SEM are shown for three experiments. **d,** Representative endonuclease assays with 100 nM hMutLγ in 5 mM Mg^2+^ or Mn^2+^ with the addition of various second metal cofactors (5 mM). Reactions were incubated at 37°C for 90 min. Metals other than Mg^2+^ compete with Mn^2+^ to inhibit hMutLγ endonuclease activity. % N, percent nicked product. Means ± SEM are shown for three experiments after subtracting background nicked product from no-protein controls. **e,** hMutLγ endonuclease activity with increasing ATP concentration. 100 nM hMutLγ was incubated with 5 mM Mn^2+^ and indicated concentrations of ATP at 37°C for 60 min. Endonuclease stimulation was optimal with 0.5 mM ATP, while higher concentrations were inhibitory.

**Extended Data Figure 2.**
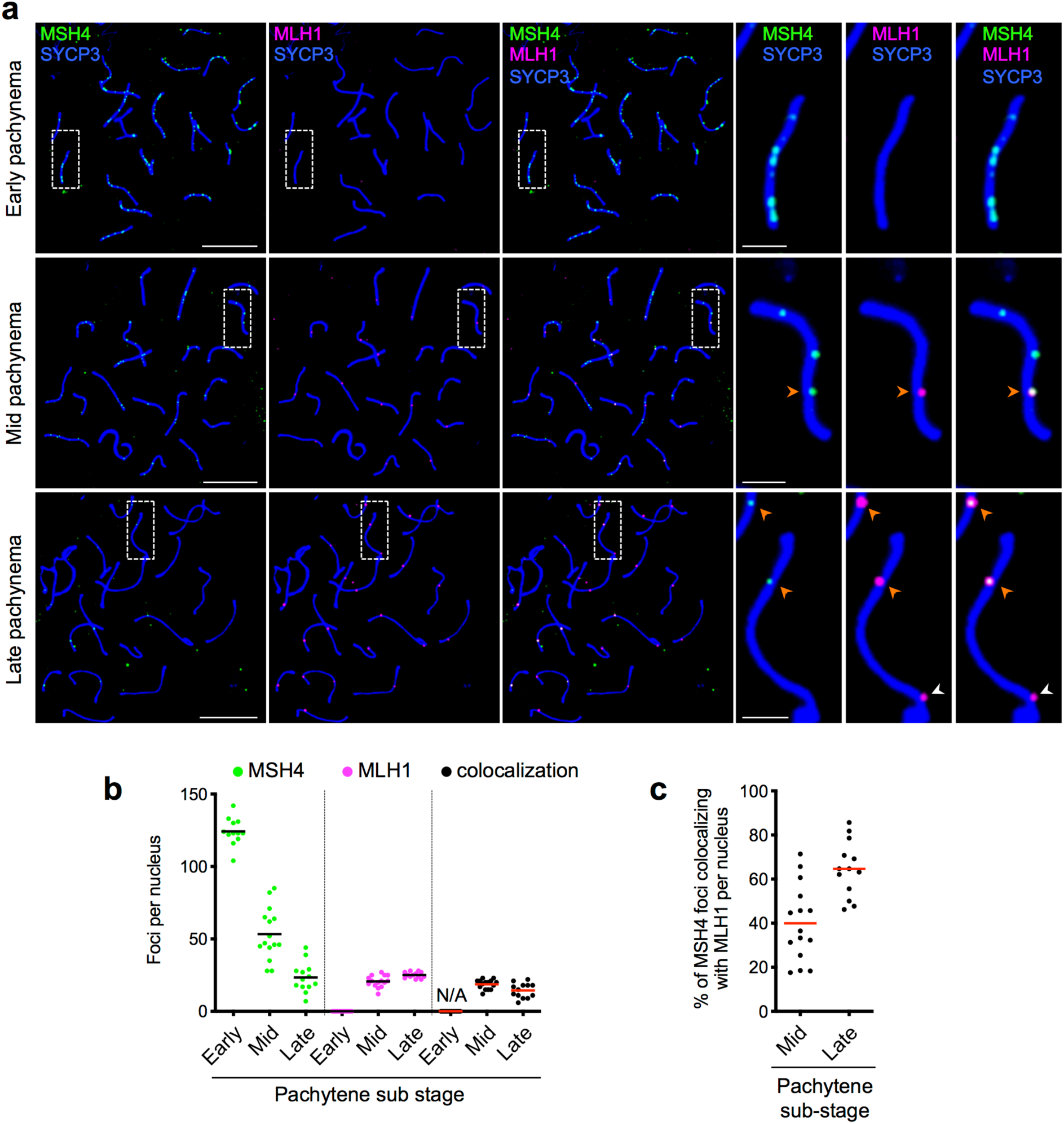
Colocalization of MutLγ and MutSγ in mouse spermatocytes. **a,** Representative images of surface-spread spermatocyte chromosomes from the indicated pachytene substages, immuno-stained for MSH4 (green), MLH1 (magenta) and the chromosome axis marker SYCP3 (blue). Magnified panels show individual pairs of synapsed chromosomes. Arrowheads indicate crossover-specific MLH1 foci that colocalize with MSH4. **b,** Quantification of MSH4 and MLH1 foci and MSH4-MLH1 cofoci in mid- and late-pachytene stages (N/A, not applicable because MLH1 foci were not observed in early pachytene). **c,** Percent of MSH4 foci that are colocalized with MLH1 in mid and late pachytene nuclei. The fraction of co-foci increases as the other MSH4 foci disappear in late pachynema.

**Extended Data Figure 3.**
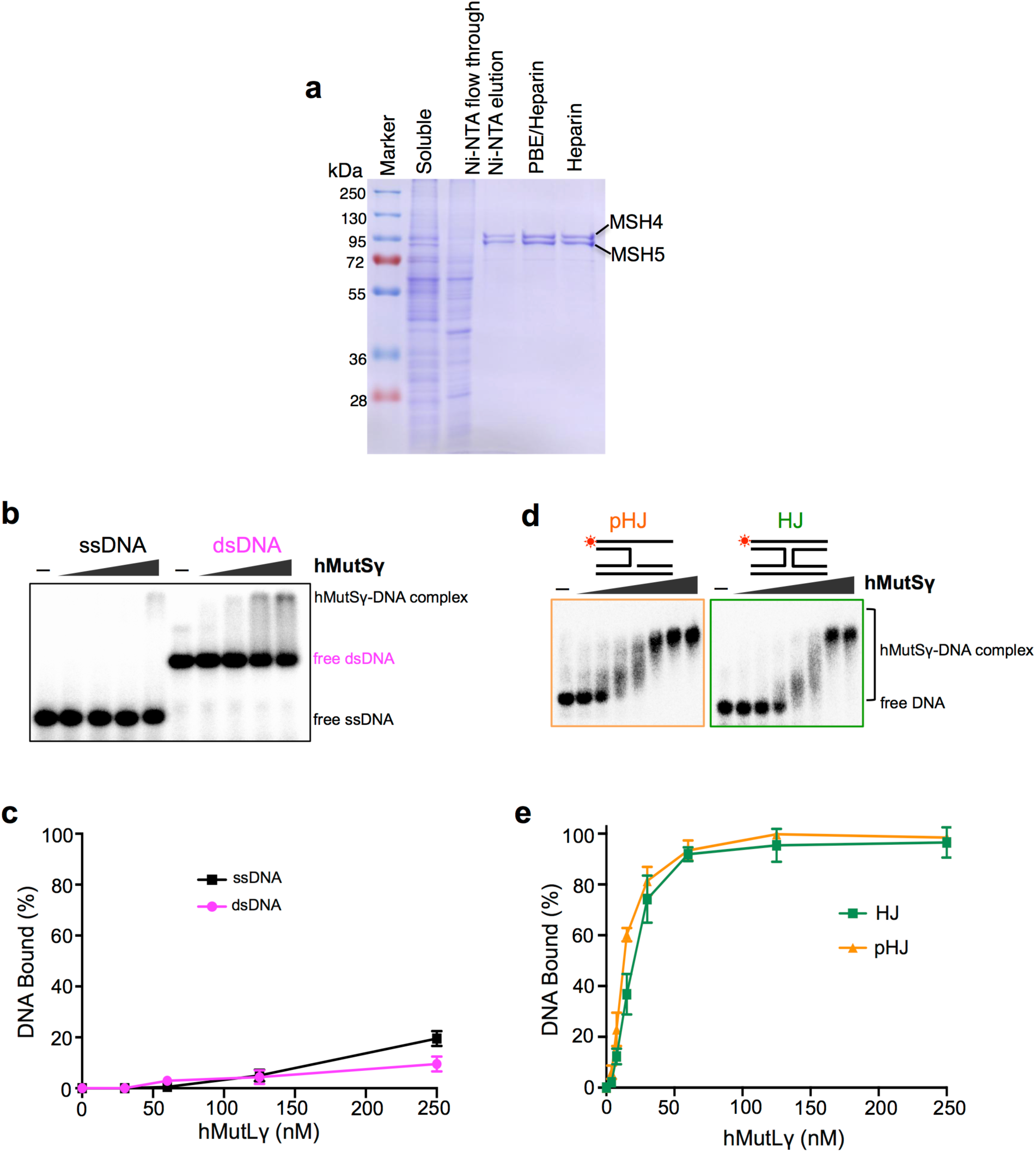
Purification and characterization of human MutSγ. **a,** Representative hMutSγ purification steps monitored by 10% SDS-PAGE stained with Coomassie blue. **b,** Representative images of electrophoretic mobility shift assays (EMSAs) analyzing hMutSγ binding to 80mer single and double-stranded DNAs. **c,** Quantification of the EMSAs represented in panel **b** showing means ± SEM for three independent experiments. **d,** Representative images of EMSAs analyzing hMutSγ binding to pro-HJ and Holliday junction structures. Terminally ^32^P-labeled strands are indicated by asterisks. **e,** Quantification of the EMSAs represented in panel **d** confirm high affinity binding of hMutSγ to pHJ and HJ structures. In **c** and **e**, means ± SEM for three independent experiments. Binding reactions contained 5 nM ^32^P-labeled DNA, 100 mM NaCl and 5 mM Mg^2+^ and were incubated on ice for 10 min. Bound and free DNA species were resolved by non-denaturing 5% polyacrylamide gel electrophoresis and processed for imaging.

**Extended Data Figure 4.**
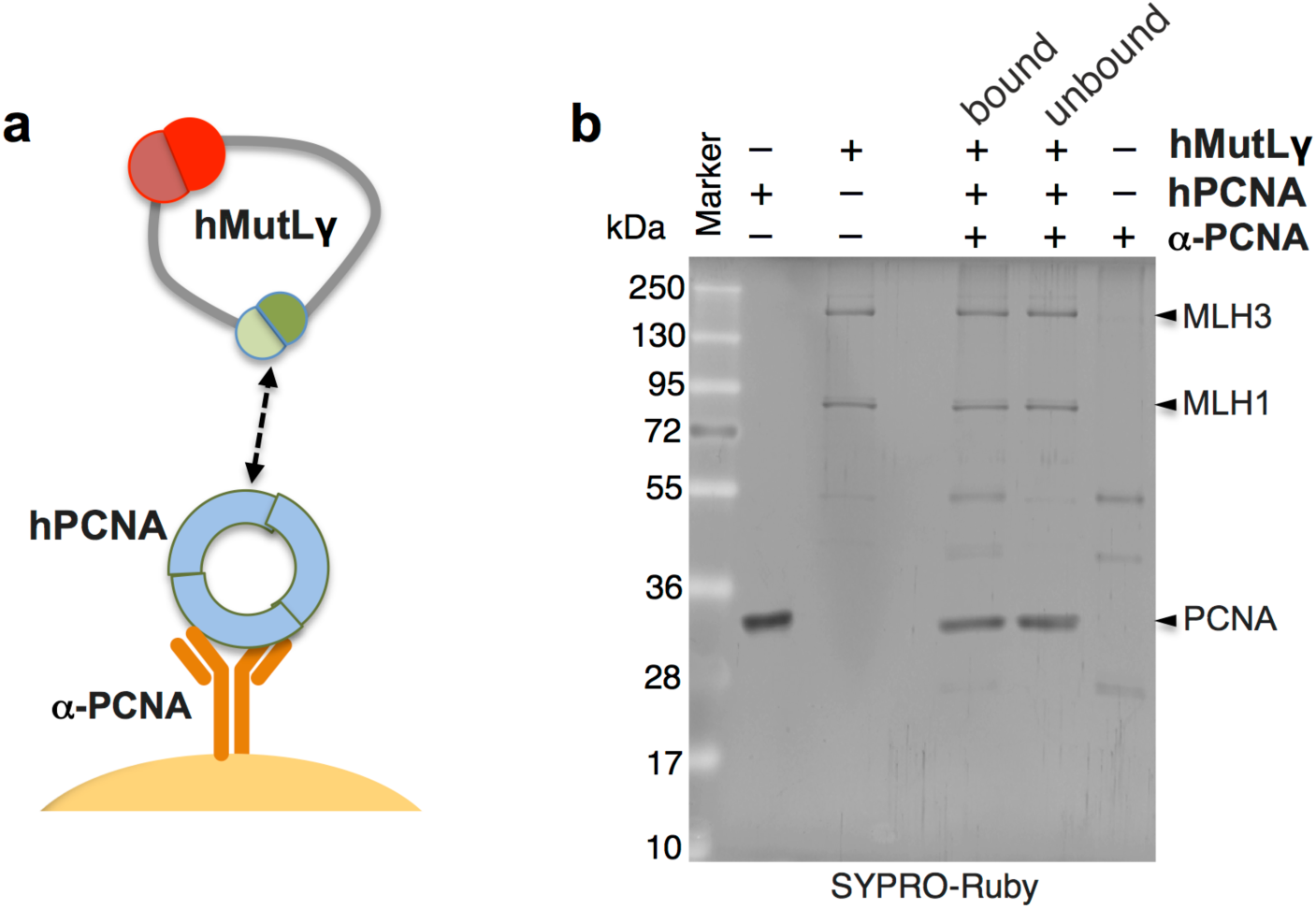
hMutLγ and hPCNA interact in solution. **a,** Experimental scheme to analyze hMutLγ-hPCNA interaction in solution. **b,** SYPRO-Ruby stained SDS-PAGE analysis of hMutLγ-hPCNA interaction.

**Extended Data Figure 5.**
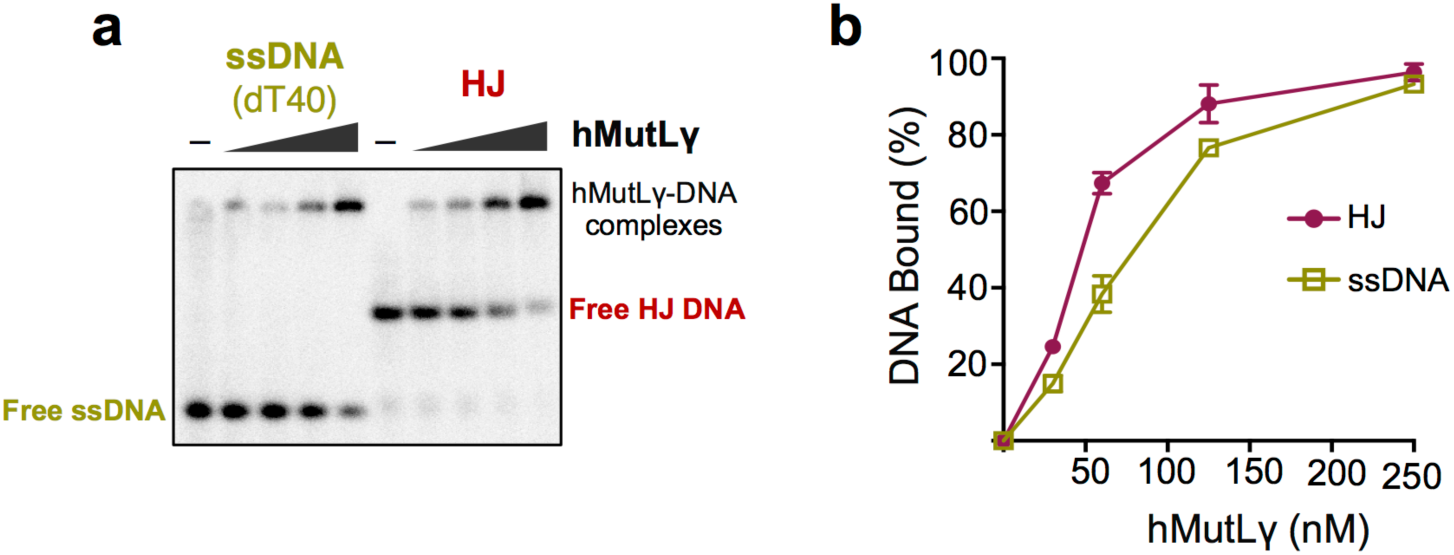
ssDNA binding by hMutLγ. **a,** Representative image of EMSAs comparing hMutLγ binding to ssDNA and Holliday junctions.. **b,** Quantification of the EMSAs represented in panel **a**. Means ± SEM for three independent experiments are shown. Binding reactions contained 1 nM ^32^P-labeled DNA, 100 mM NaCl and 5 mM Mg^2+^ and were incubated on ice for 10 min. Bound and free DNA species were resolved by non-denaturing 5% polyacrylamide gel electrophoresis and processed for imaging.

**Extended Data Figure 6.**
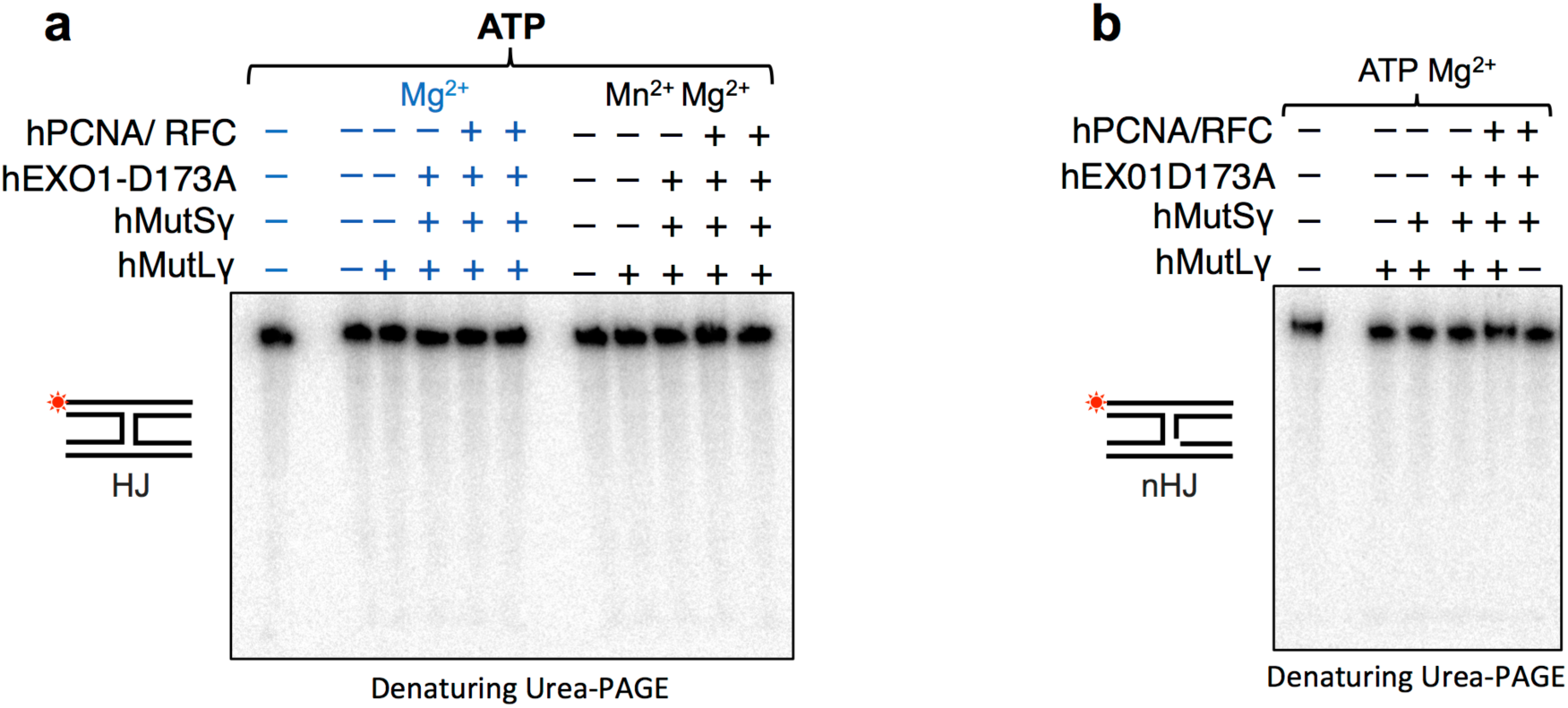
Model Holliday junctions are not incised by hMutLγ. **a** and **b,** Representative gel images of ensemble endonuclease reactions with HJ and nicked HJ substrates. Reactions contained 25 nM indicated proteins, 0.5 mM ATP and 5 mM Mg^2+^/Mn^2+^, and were incubated at 37°C for 30 min. The reaction products were analyzed on a 15% denaturing polyacrylamide gel to detect nicking at any position. Nicked DNA strands were not detected and non-denaturing gels showed that model HJs and nHJs were not resolved by the hMutLγ ensemble (not shown).

**Extended Data Figure 7.**
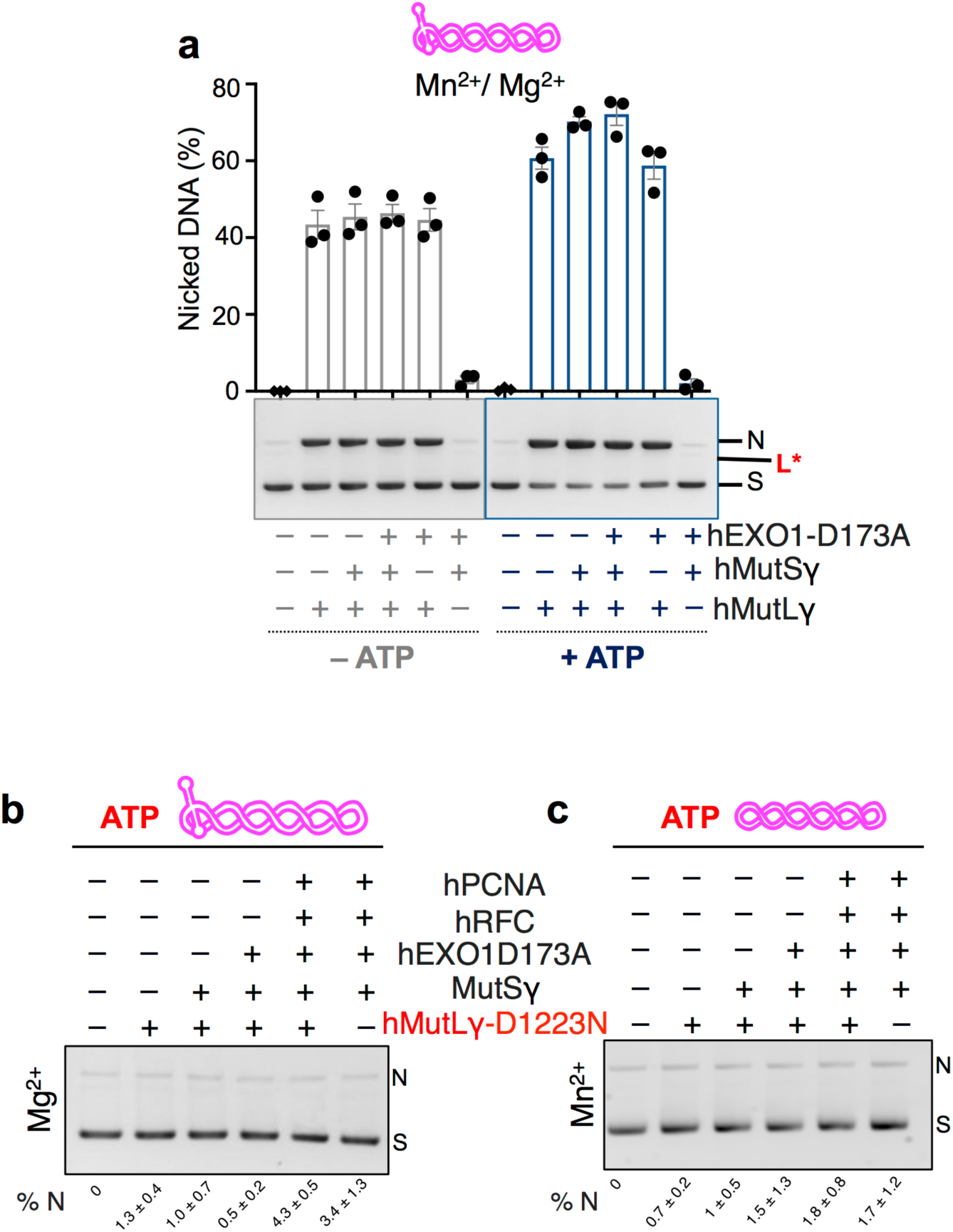
Endonuclease assays the HJ-containing pUC(AT) DNA substrate on hMutLγ ensemble endonuclease reactions. **a,** Endonuclease assays with pUC(AT) and varying mixtures of hMutLγ, hMutSγ and hEXO1-D173A, with and without ATP (50 nM each protein, 0.5 mM ATP, 5 mM Mn^2+^ and 5 mM Mg^2+^, incubated at 37°C for 60 min). Means ± SEM are shown for three experiments. Under these conditions, pUC(AT) is incised 1.3-1.7-fold more efficiently than the corresponding reactions with pUC19 (see Fig. 2g). Notably, the linear cleavage product (L*) seen with pUC19 (Fig. 2g) was not observed with pUC(AT). Thus, both the efficiency of nicking and the nature of the incision products are altered when a Holliday junction is present. **b** and **c,** Negative control ensemble endonuclease assays for pUC(AT) (**b**) and pUC19 (**c**) containing nuclease-dead hMutLγ-D1223N. Reactions contained 25 nM indicated proteins, 0.5 mM ATP and 5 mM Mg^2+^ or Mn^2+^ and were incubated at 37°C for 30 min. Average percent nicking (%N) ± SEM are shown for three experiments.

**Extended Data Figure 8.**
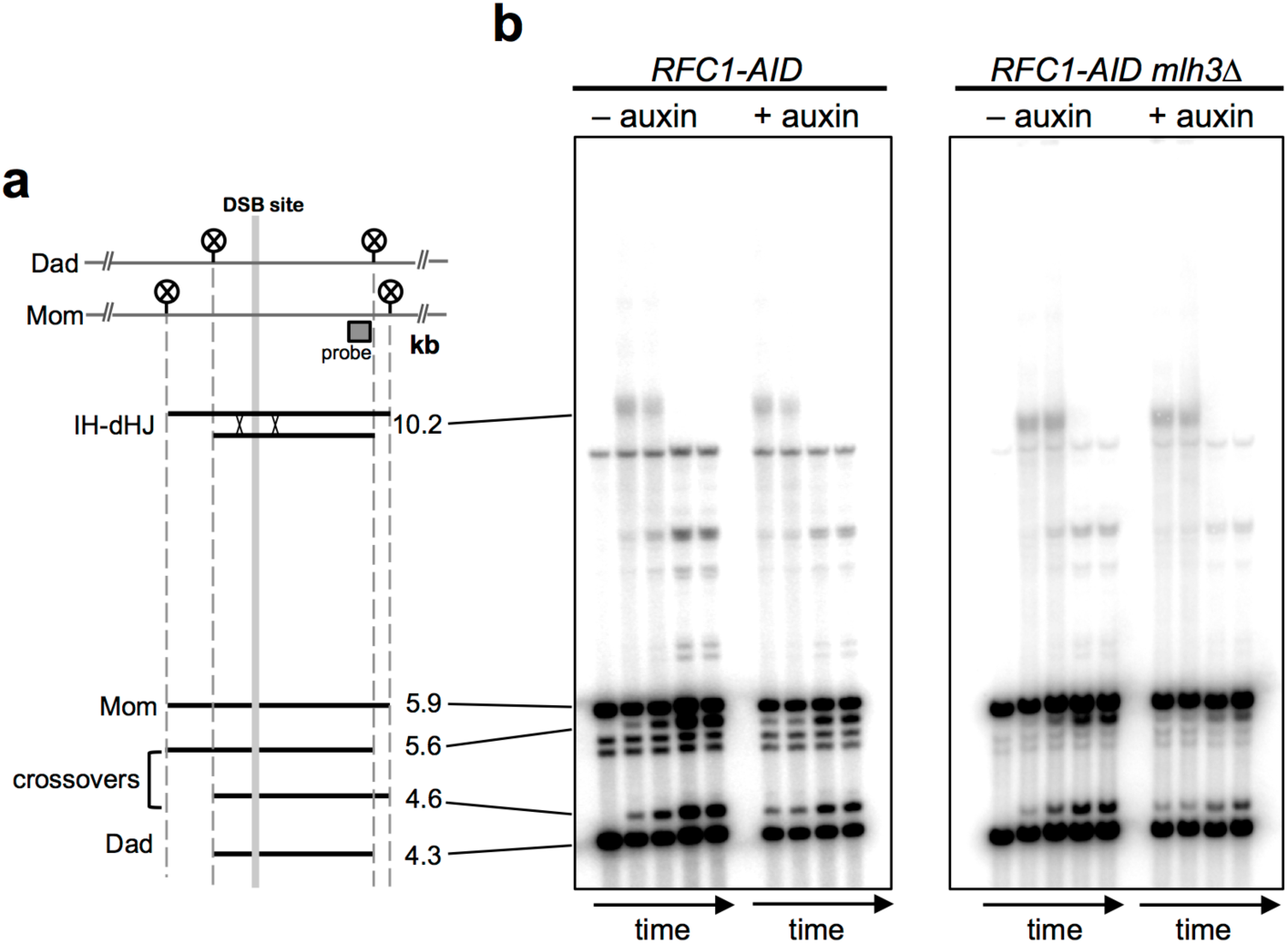
Physical assay system for analysis of meiotic recombination n budding yeast. **a,** Map of the *HIS4:LEU2* recombination hotspot locus highlighting the initiating DSB site, *Xho*I restriction sites (circled Xs) and the position of the probe used in Southern blotting. Sizes of diagnostic fragments are shown below. **b,** Representative 1D gel Southern blot images for analysis of crossovers and dHJs in *RFC1-AID* and *RFC1-AID mlh3Δ* strains, with and without the addition of auxin to trigger Rfc1-AID degradation. Time points are 0, 7, 8, 9, and 11 hours after induction of meiosis.

## Supplementary Data

**Table S1.**
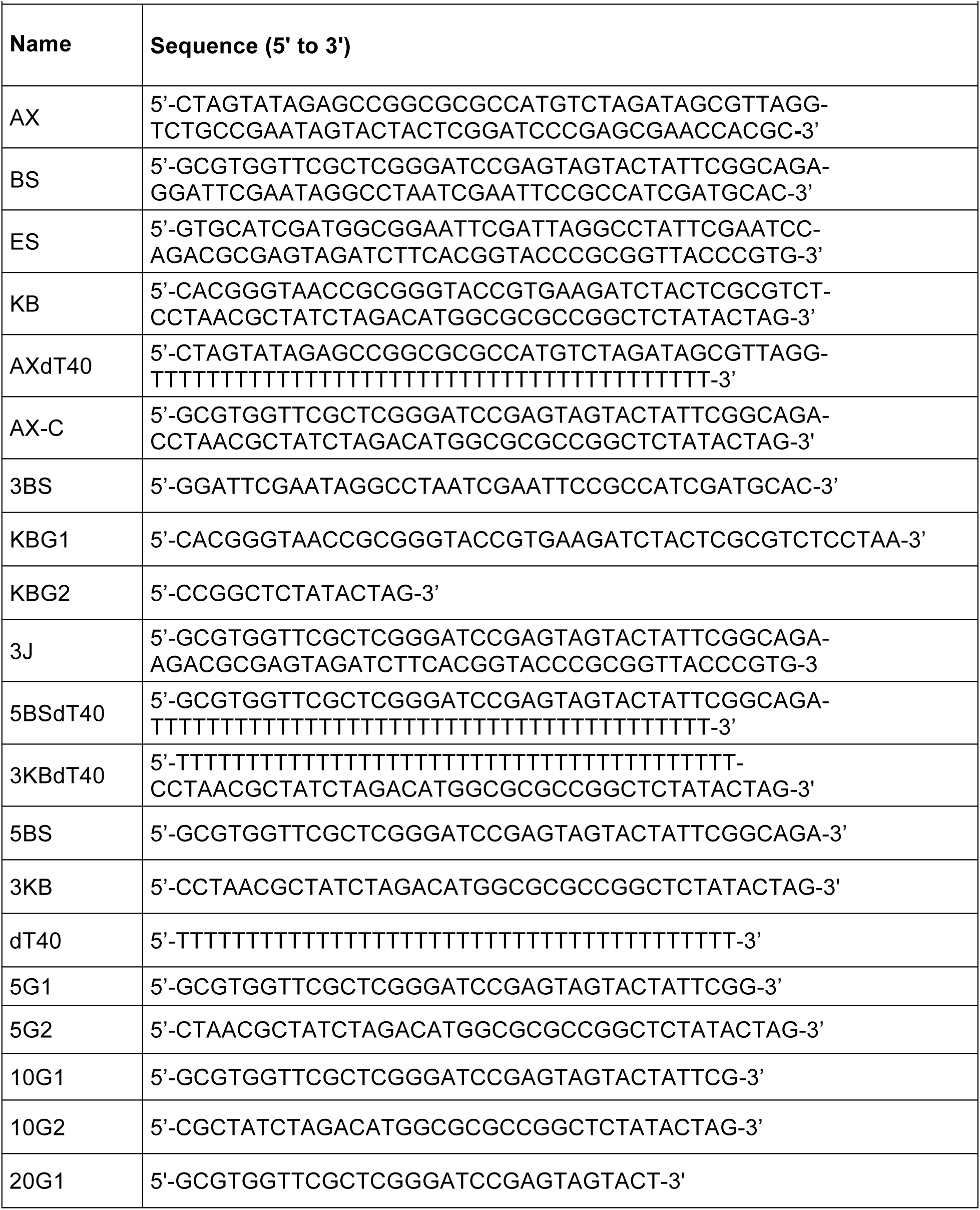
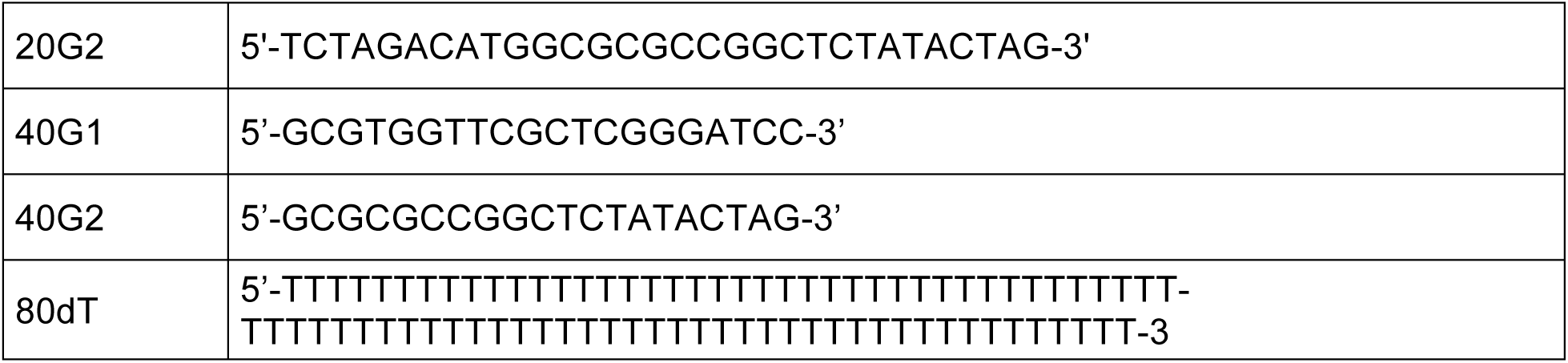
Oligonucleotides used in this study.

**Table S2.**
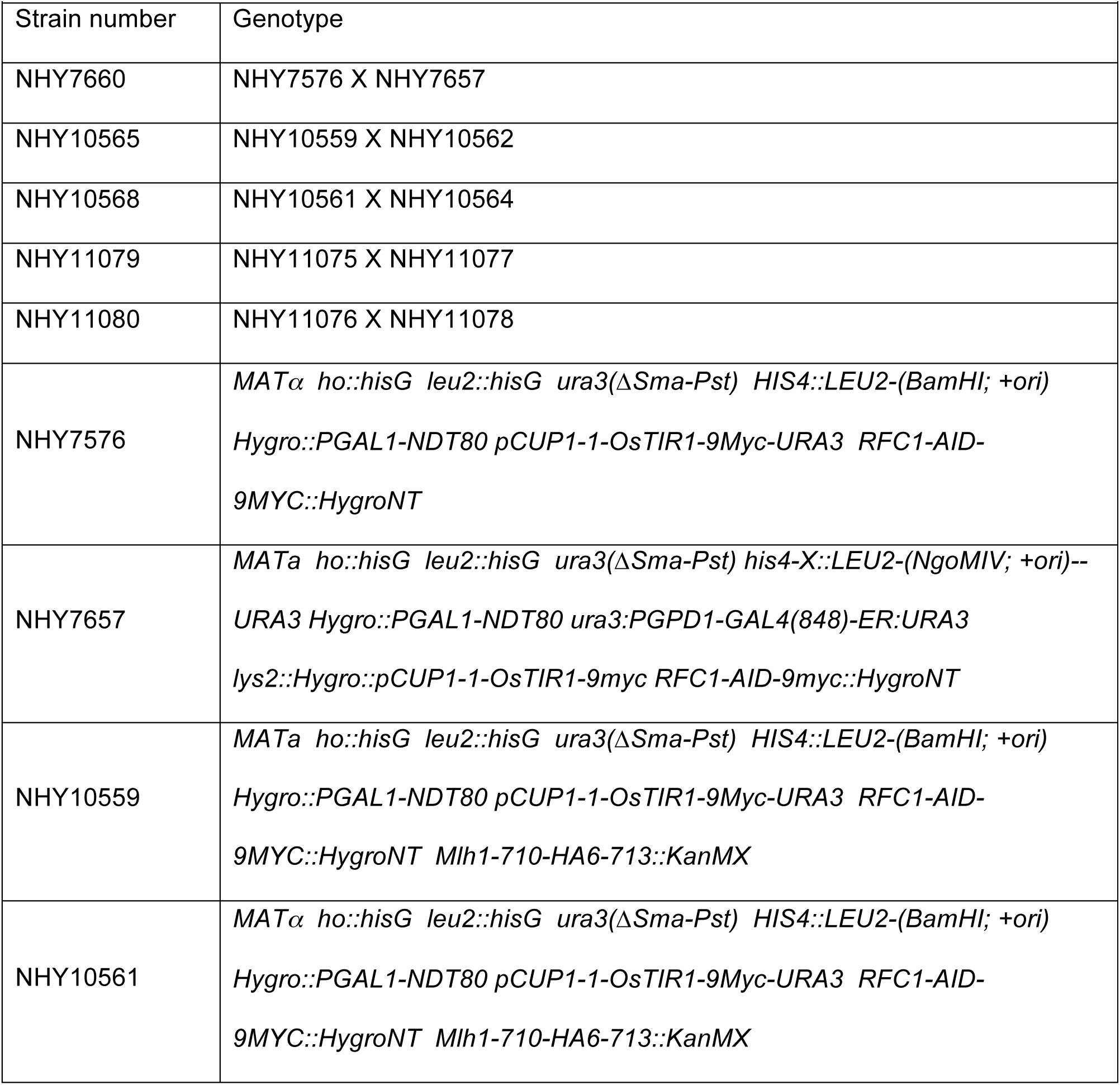
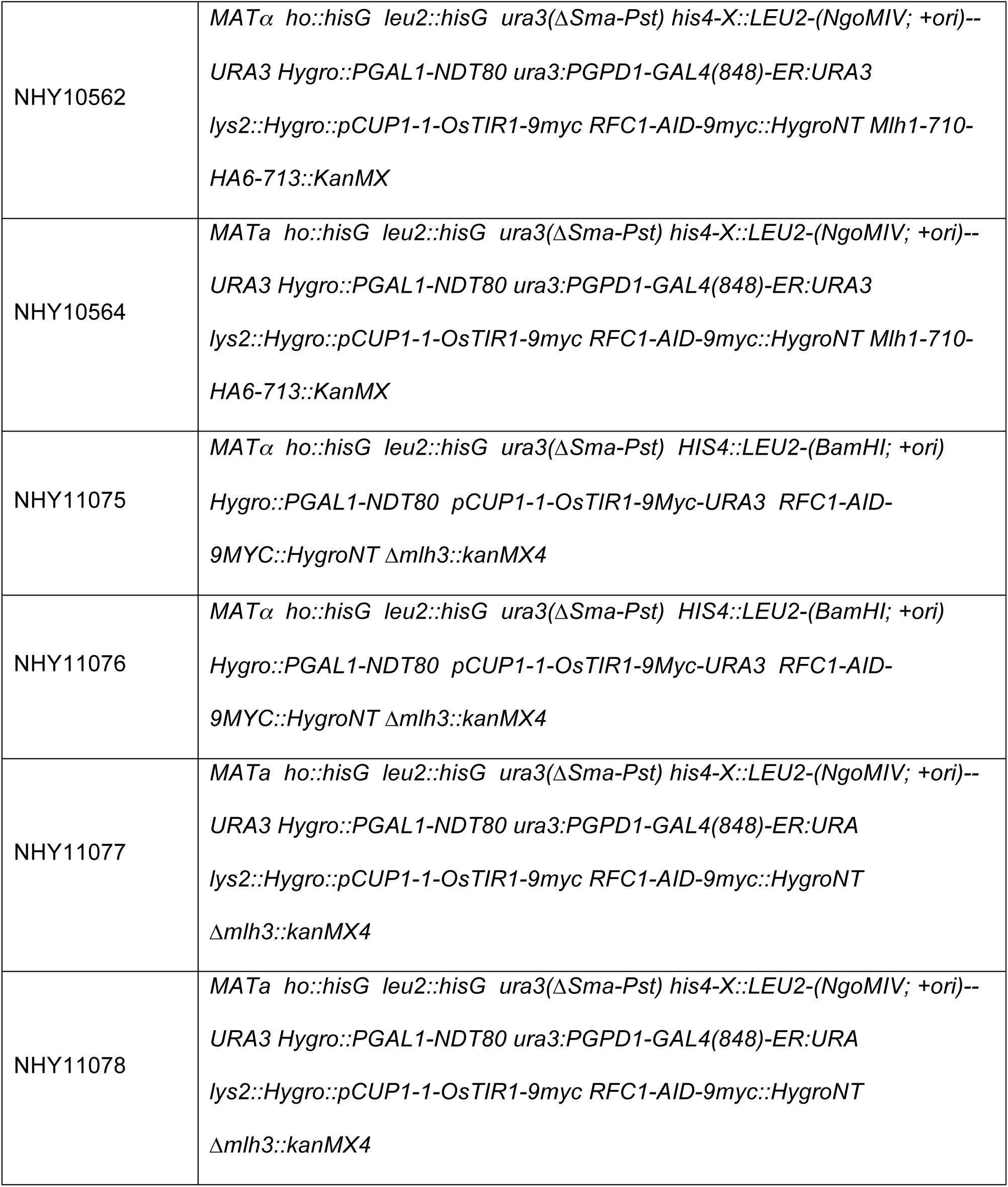
Strain used in this study

